# Cell-free expression with a quartz crystal microbalance enables rapid, dynamic, and label-free characterization of membrane-interacting proteins

**DOI:** 10.1101/2024.01.14.575553

**Authors:** Aset Khakimzhan, Ziane Izri, Seth Thompson, Oleg Dmytrenko, Patrick Fischer, Chase Beisel, Vincent Noireaux

## Abstract

Integral and interacting membrane proteins (IIMPs) constitute a vast family of biomolecules that perform essential functions in all forms of life. However, characterizing their interactions with lipid bilayers remains limited due to challenges in purifying and reconstituting IIMPs *in vitro* or labeling IIMPs without disrupting their function *in vivo*. Here, we report TXTL-QCMD to dynamically characterize interactions between diverse IIMPs and membranes without protein purification or labeling. As part of TXTL-QCMD, IIMPs are synthesized using cell-free transcription-translation (TXTL), and their interactions with supported lipid bilayers are measured using a quartz crystal microbalance with dissipation (QCMD). TXTL-QCMD reconstitutes known IIMP-membrane dependencies, including specific association with prokaryotic or eukaryotic membranes or oscillating interactions by the *E. coli* Min system. Applying TXTL-QCMD to the recently discovered Zorya anti-phage system unamenable to labeling, we discovered that ZorA and ZorB integrate together within the lipids found at the poles of bacteria while ZorE diffuses freely on the non-pole membrane. These efforts establish the potential of TXTL-QCMD to broadly characterize the large diversity of IIMPs.

---

Lipid bilayers form the physical boundaries between the inner compartment of a living cell and the environment. To sense and interact with their surroundings, cells synthesize integral and interacting membrane proteins (IIMPs) that localize either into or at the surface of phospholipid membranes. IIMPs constitute a large family of biomolecules achieving broad cellular functions that interface cells with their milieu^1,2^. Lipid bilayers also serve as physical templates for IIMPs to organize cellular functions, comprising, for instance, the formation of dynamical patterns^3^ and anchoring cytoskeletons^4,5^. Characterizing IIMPs non-disruptively, such as the biochemistry and biophysics of their interactions with membranes, is often challenging to achieve *in vivo* as they usually require labeling with fluorescent reporters or affinity tags. Conversely, the reconstitution of IIMPs *in vitro* provides easier access to their biochemical and biophysical characterization. Yet, this approach necessitates difficult recombinant purification and reconstitution procedures that prevent rapid exploration of their properties.

Cell-free transcription-translation (TXTL) simplifies and outpaces traditional recombinant approaches by enabling, in a matter of hours, the scalable synthesis of proteins outside living cells, including IIMPs, from plasmids or linear DNA^6–8^. Many TXTL reactions can be prepared concurrently, thus facilitating the rapid and parallel characterization of reaction products. To fold properly and not precipitate, IIMPs synthesized in TXTL require either the presence of non-natural substrates like surfactants or natural membranes made of phospholipids^9–12^, which is typically achieved by adding liposomes or nanodiscs to the TXTL reaction. Liposomes are known to be unstable in TXTL reactions^13^, whereas nanodiscs are soluble lipid rafts stable in cell-free reactions^13–15^. While nanodiscs offer a simple method to synthesize IIMPs in TXTL, they are limited in size (10-20 nm), preventing the formation of large IIMP complexes, and lipid compositions^16^, reducing the scope of IIMP-lipids interactions that can be studied. Characterizing the interaction with or the integration of IIMPs into nanodiscs requires either disruptive fluorescent labeling or techniques such as gel electrophoresis^17^. IIMPs can also be synthesized in TXTL reactions encapsulated into liposomes, a system known as synthetic cells^15,18,19^. In these settings too, disruptive fluorescent labeling is required to visualize the interaction with the membrane. Preparing TXTL-based synthetic cells with complex membrane compositions, such as *E. coli* phospholipids, is limited, as the efficacy of liposome production rapidly decreases with the complexity of the lipid composition. An approach enabling the sensitive and label-free characterization of IIMPs’ interactions with lipid membranes of arbitrarily complex membrane composition and on unlimited physical scales is still lacking.

In this work we achieved rapid, sensitive, and label-free characterization of IIMPs’ interactions with supported lipid bilayers (SLBs) by combining the versatility of TXTL with the sensitivity of a quartz crystal microbalance with dissipation (QCMD)^20^ **(Figure 1)**. We use the change in frequency of the QCMD, which is proportional to the mass added to the SLB, to measure the interaction between IIMPs and a given phospholipid composition of the bilayer. The proof of concept of such an approach has been established for single IIMPs^21,22^, but the method has never been generalized to multiple and interacting IIMPs, membrane-dependent dynamical patterns, and complex phospholipid membrane compositions. This approach provides a high signal-to-noise ratio due to minimal nonspecific adsorption of the TXTL components onto SLBs. IIMPs, without any modifications such as the addition of fluorescent reporters, are dynamically synthesized on the SLBs inside the QCMD while monitoring the signal, which eliminates recombinant expression, purification, and reconstitution procedures. SLBs made of complex phospholipid compositions, analogous to the ones found in living cells, can be prepared onto the QCMD chips, circumventing the limitations of the other methods.

**Figure 1.**
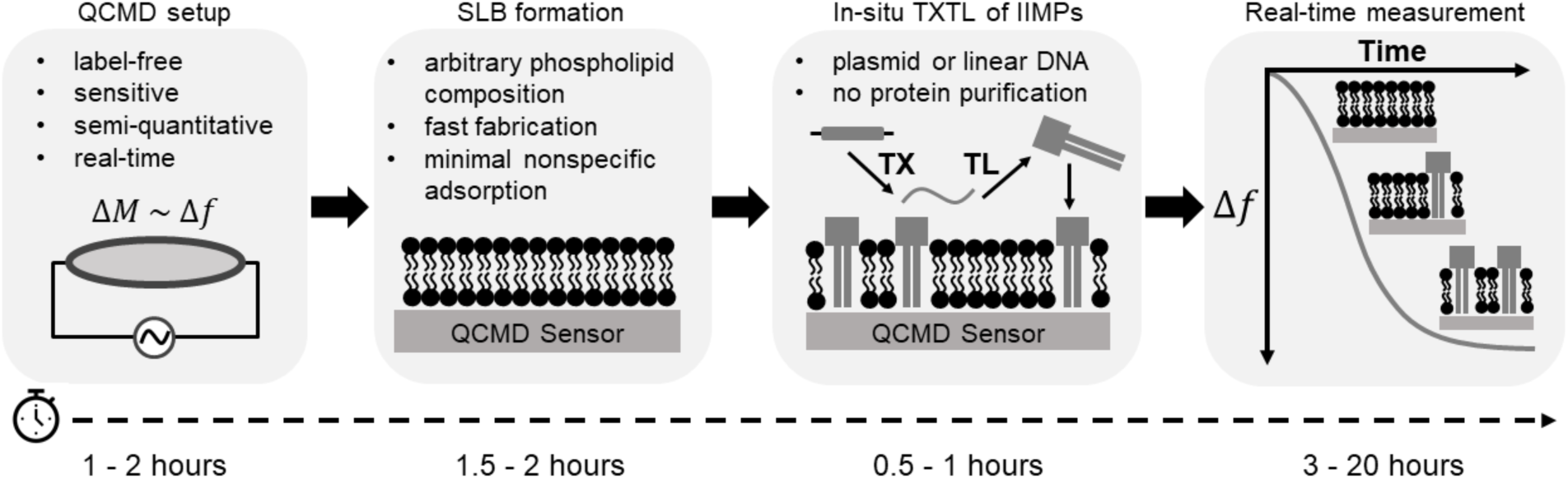
Workflow of the TXTL-QCMD experimental procedure. The TXTL-QCMD procedure allows non-disruptive semi-quantitative measurement of the interactions between *in situ* synthesized proteins and lipid bilayers. The first step of the procedure is the QCMD module setup, which takes approximately two hours. The second step of the experiment is the fabrication of the supported lipid bilayer (SLB). Using the solvent-assisted lipid bilayer formation method (SALB), SLBs of arbitrary phospholipid compositions are made in 2 hours. The third step is the preparation of the TXTL reaction, which involves mixing the TXTL components with a plasmid or a linear construct of interest. TXTL enables characterizing IIMPs that require co-translational integration. The last step is the incubation of the prepared TXTL reactions in contact with the SLB-sensor system. Typically, the reactions are incubated overnight; however, in some cases, as few as 3 hours are sufficient to obtain experimentally relevant data.

First, we characterize the nonspecific adsorption of blank TXTL reactions on the SLBs. Next, we use Alpha-Hemolysin (AH) from *S. aureus* and the large mechano-sensitive channel (MscL) from *E. coli* as model IIMPs to test the specificity of their integration into a eukaryotic membrane made of phosphatidylcholine (PC) and a bacterial membrane made of *E. coli* lipids (ECL). We observe that AH preferentially integrates into the PC membrane, whereas MscL preferentially integrates into the ECL membrane. We create a hybrid membrane composed of PC and ECL capable of hosting both MscL and AH. Binding of the C2-domain of lactadherin (Lact-C2) to the surface of lipid bilayers is observed specifically when phosphatidylserine (PS)^23^ is added to membranes, which demonstrates that proteins binding to the surface of membranes can also be sensed. We show that the *E. coli* Min system, composed of the proteins MinD, MinE, and MinC, produces oscillations on the 1.2-cm diameter QCMD sensor. Oscillations are observed for broad varieties of lipid bilayer compositions and synthesized Min proteins. Surprisingly, MinD alone, without MinE, produces oscillations. Lastly, we apply our method to study the previously uncharacterized microbial anti-phage system, Zorya, hypothesized to be membrane-dependent^24^. We confirm this hypothesis and further suggest potential mechanisms for Zorya-based immunity. QCMD combined with TXTL turns out to be a fast analytical method to obtain information on IIMPs’ interactions with lipid membranes that is useful for basic biology discoveries and for the intermediate bioengineering of systems integrating IIMPs, such as synthetic cell systems^19^.

## Results and Discussion

### TXTL reactions have minimal nonspecific adsorption on SLBs

The TXTL system, known as myTXTL (Daicel Arbor Biosciences), uses the endogenous *E. coli* core RNA polymerase and sigma factor 70 present in the lysate as the sole primary transcription mechanism. This system does not contain any remaining live *E. coli* cells **(Figure S1)**. Genes are expressed either from plasmids or linear dsDNA, from *E. coli* promoters (*e.g.*, P70a), or from the T7 promoter via a transcriptional activation cascade **(Figure S2)**, as previously described^17,25,26^.

The interaction with and integration of IIMPs into membranes depend on the membrane’s phospholipid composition^27^. Consequently, the diversity of SLBs that can be fabricated on the QCMD sensor determines the breadth of IIMPs that can be characterized, in particular IIMPs specific to the membranes of either bacteria or higher organisms. To make SLBs on the QCMD sensor, we used the Solvent Assisted Lipid Bilayer Formation Method (SALB)^28–30^, as it is fast and consistent **(Figure 2a)**. Our model eukaryotic membrane is made of EggPC, a mixture of PC phospholipids extracted from chicken eggs, with various aliphatic chain lengths. Our model prokaryotic cell membrane is made of ECL that consists of a mixture of phospholipids extracted from *E. coli* composed of mainly PE (phosphatidylethanolamine, ∼75% mol), PG (phosphatidylglycerol, ∼20% mol), and CL (cardiolipin, ∼5% mol)^31^. We studied interactions of IIMPs with many other SLBs made from mixtures of pure phospholipids. In these cases, the SLBs included the most abundant phospholipid found in EggPC (DOPC) or ECL (DOPE). A TXTL reaction contains a broad variety of biological molecules in large concentrations; it is thus important to make sure that they do not produce large nonspecific adsorption on the sensor-SLB system in the absence of IIMPs. We devised a procedure so that SLBs cover the whole sensor and have no nonspecific adsorption.

**Figure 2.**
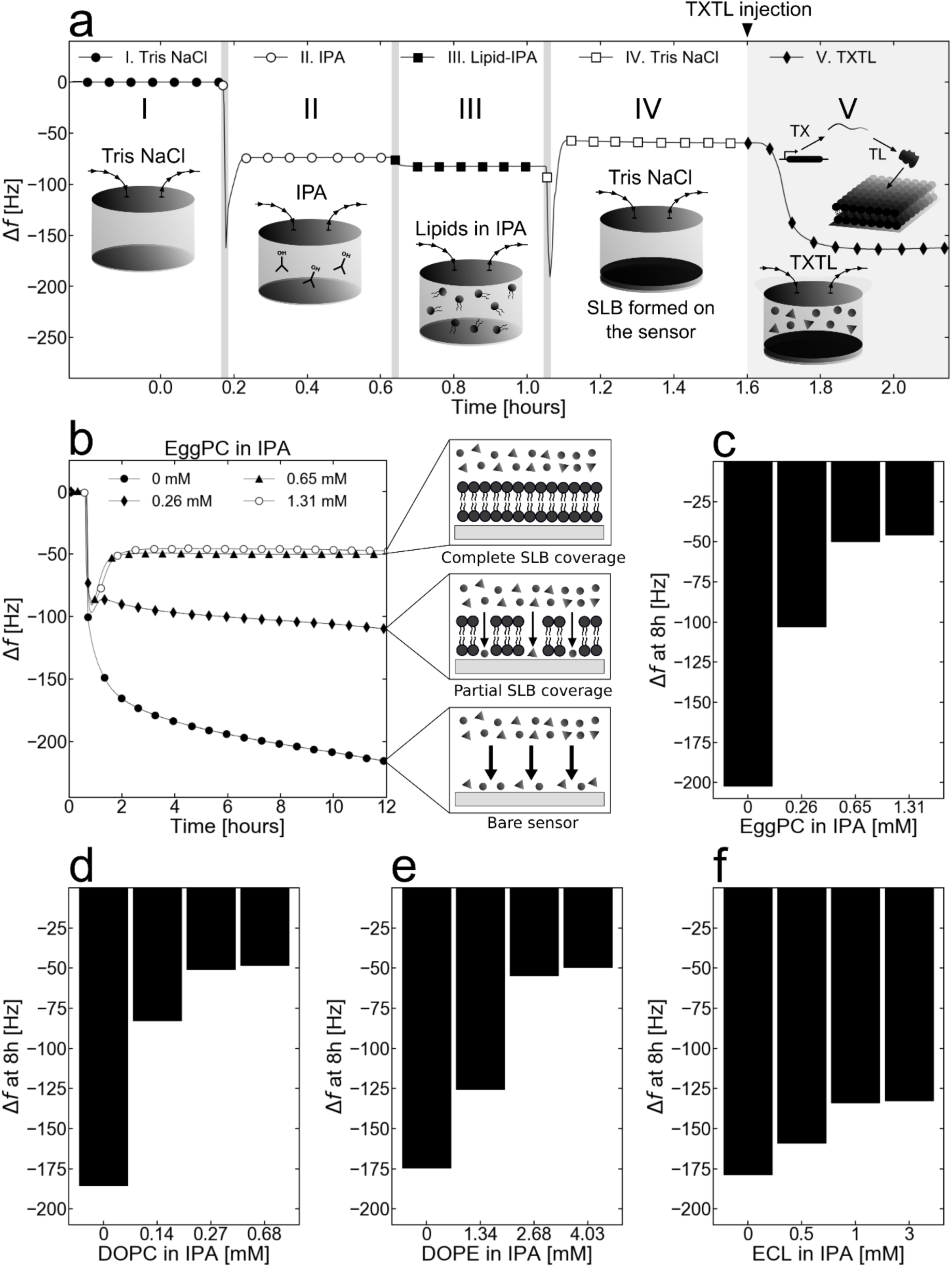
An optimal lipid-isopropanol concentration during SALB is critical for complete SLB coverage and minimal TXTL nonspecific adsorption. **a.** Experimental steps of SALB formation (steps I-IV) followed by the nonspecific adsorption characterization with a TXTL reaction that only synthesizes the T7 RNA polymerase (step IV, plasmid P70a-*T7rnap* 0.15 nM). During step I, the sensor is prepared for SLB formation measurements by equilibrating the QCMD signal with a flush of the Tris-NaCl buffer. Step II of SALB formation is the flushing of isopropanol (IPA) to remove any residual Tris NaCl. The dip in the frequency followed by the stabilized 70-80 Hz frequency shift in this step is attributed to the change of the liquid from the aqueous Tris-NaCl buffer to IPA. Step III is the flushing of the phospholipid-IPA solution. Here, because the phospholipids are added to IPA, the frequency changes by 5-10Hz, which indicates a change in the characteristics of the flushed liquid. Step IV is the second Tris NaCl buffer flush, during which the SLB is formed. The bilayer forming mechanism of SALB relies on the gradual change of the liquid in the module from IPA to an aqueous buffer. The mass and properties of the SLB are reflected in the frequency difference between step IV and the initial Tris NaCl flush of step I. In step V, the TXTL reaction is flushed, and the incubation begins. **b.** The adsorption kinetics of a blank TXTL reaction (plasmid P70a-*T7rnap*, 0.15 nM) depending on the EggPC phospholipids concentration in IPA during step II. The cartoon frames on the right explain the different levels of non-specific adsorption to the SLB-sensor system. **c.** Changes in frequency after 8 hours of incubation (blank TXTL reaction with only P70a-*T7rnap*, 0.15 nM) as a function of phospholipids concentration in IPA during step II for EggPC. **d., e., and f.** Same as **c.** for DOPC, DOPE, and *E. coli* lipids (ECL) total respectively.

The SLBs were made in four steps **(Figure 2a)**. In step I, the QCMD sensor is washed with a Tris NaCl buffer, which is replaced in step II by an isopropyl alcohol (IPA) solution. In step III, phospholipids dissolved in IPA are introduced onto the sensor. The spontaneous SLB formation occurs in step IV, as the IPA with the dissolved phospholipids is gradually replaced with a Tris NaCl buffer^30^. During these steps, the resonance frequency of the SiO_2_ crystal of the sensor varies: the more mass is adsorbed to it, the smaller the resonance frequency^32^. The difference between the stabilized resonance frequency (Δf = 0) from steps I to IV is negative and increases in absolute value with the mass of the SLB on the sensor^33,34^. When no phospholipids are added to the IPA during step III, the difference is zero (**Figure S3**). In all the subsequent data presented, the resonance frequency at the end of step IV, before adding the TXTL reaction onto the QCMD sensor, was taken as the reference and set to zero. A short-lived dip in frequency is systematically produced at the beginning of each step. This is due to the introduction of a different solution into the QCMD modules **(Figure 2a)**. This short-lived dip in frequency, also present when a TXTL reaction is introduced into the QCMD chambers, is irrelevant to the measurement of the interaction between the IIMPs and the SLBs.

To determine whether an SLB fully covers the sensor, is stable, and does not allow nonspecific TXTL adsorption, a blank TXTL reaction (with only the plasmid P70a-*T7rnap*, 0.15 nM) was incubated into the QCMD module in contact with the SLB-sensor system. SLBs were made with different concentrations of phospholipids to determine at which phospholipid concentration nonspecific TXTL adsorption is not observed. This assay was carried out for each SLB tested in this work. In the case of an EggPC membrane, the full coverage of the QCMD sensor is achieved at a phospholipid concentration at or above 0.65 mM (0.5 mg/ml) in IPA, as observed by the flat frequency shifts **(Figure 2b)**. Below a concentration of 0.65 mM, an EggPC membrane shows nonspecific TXTL adsorption as observed by the decrease in Δf for lower phospholipid concentration **(Figure 2c)**. The minimal phospholipid concentration to get SLBs that fully cover the QCMD sensor was first determined for our four base SLBs, namely EggPC, DOPC, DOPE, and ECL SLBs (**Figure 2c, d, e, f**, **Figure S4, Table S1)**. We found that the four major phospholipid compositions used in this work can be used reproducibly for QCMD-TXTL experiments without nonspecific adsorption from the TXTL components **(Table S1)**. Achieving full coverage of the QCMD sensor with lipid bilayers is a striking result that shows the natural resistance of phospholipid membranes to nonspecific adsorption of a complex physiological solution like a TXTL reaction composed of 10 mg/ml of proteins, tRNAs, and rRNAs, about 250-300 mM salts and many other chemicals^26,35^.

### Cell-free synthesized IIMPs interact specifically with their natural membranes

To test the QCMD-TXTL approach to membrane protein-lipid interaction specificities, we chose two proteins known to preferentially reside in different natural membranes. We performed a set of TXTL reactions that synthesize either the pore-forming protein Alpha-Hemolysin (AH) from *S. aureus*, known to target mammalian cells rich in PC^36^, and the large-conductance mechanosensitive channel MscL from *E. coli* in membranes (ECL) mostly composed of PE, PG, and CL. To quantify the amount of protein produced in the QCMD chamber, we used a fusion AH-eGFP proven to be functional in TXTL because it is produced as a soluble protein and thus easy to quantify^37^. Both genes were cloned under the T7p14 promoter and expressed via the T7 transcriptional activation cascade (T7p14-*ah-egfp*, T7p14-*mscL*) (**Figure 3a**). AH assembles into a heptameric pore^38^, while MscL assembles into a pentamer channel^39^. We observed a net frequency drop only when MscL was synthesized onto the ECL SLB. Conversely, we observed a net frequency drop only when AH-eGFP was synthesized onto the EggPC SLB (**Figure 3b, c**). Replicates of these experiments confirmed these results (**Figure S5, S6**). The same protein-lipid specificities were observed with DOPE (interaction with AH-eGFP, no interaction with MscL) and DOPC (no interaction with AH-eGFP, interaction with MscL) SLBs (**Figure S7**). A delay in frequency shift was observed for AH, presumably because it must reach a critical concentration before interacting with the membrane^40^. MscL pre-synthesized in TXTL reactions without membranes did not produce a frequency change (**Figure S8**), which is expected because its integration is coupled to its translation. Unlike MscL, AH is a soluble pore-forming toxin that can be pre-synthesized 24 hours before being added onto the QCMD chip and still integrates into the PC SLB (**Figure S9**), without a delay as it is introduced into the QCMD module at a large concentration. The concentration of IIMP produced in the QCMD chamber of volume 40 µl was estimated with AH-eGFP to be 1-2 μM **(Figure S10)**. We found this concentration range relevant as it corresponds to the average protein concentration in *E. coli*^41^. Using MscL, we estimated the resolution of the mass added to the SLB to be on the order of 1 ng, which is a hundred times more sensitive than a standard protein gel **(Figure S11)**. This estimation also shows that about twenty times more proteins are synthesized in the 40 µL-QCMD chambers with respect to the membrane’s maximum binding capacity **(Figure S11)**.

**Figure 3.**
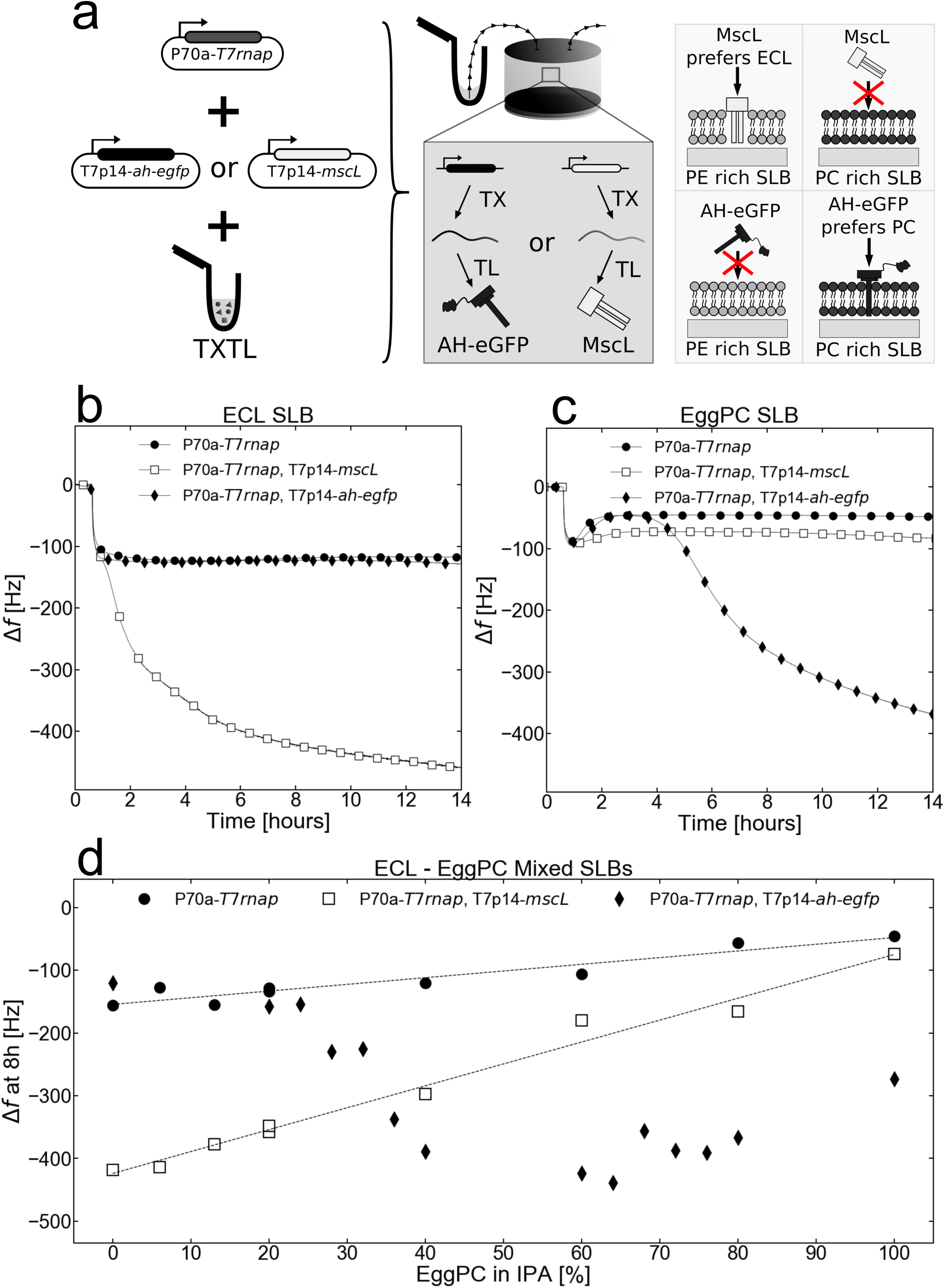
*In situ* cell-free synthesized MscL and AH-eGFP exhibit their natural phospholipid affinities in ‘pure’ and ‘mixed’ SLBs. **a.** MscL or AH-eGFP is synthesized into the QCMD module on top of either a PC-rich EggPC SLB or a PE-rich *E. coli* lipids (ECL) SLB. The rightmost cartoon demonstrates the expected protein-lipid preferences of both proteins. **b. and c.** Adsorption kinetics of a blank (P70a-*T7rnap*, 0.15 nM), an AH-eGFP (P70a-*T7rnap*, 0.15 nM, T7p14-*ah-egfp*, 5 nM), and a MscL (P70a-*T7rnap*, 0.15 nM, T7p14-*mscL*, 5 nM) synthesizing TXTL reactions in contact with an ECL SLB and an EggPC SLB respectively. **d.** The frequency changes after 8 hours of incubating either blank (P70a-*T7rnap*, 0.15 nM), AH-eGFP (P70a-*T7rnap*, 0.15 nM, T7p14-*ah-egfp*, 5 nM), or MscL (P70a-*T7rnap*, 0.15 nM, T7p14-*mscL*, 5 nM) TXTL reactions in ECL – EggPC Mixed SLBs. The x-axis percentage indicates the concentration of EggPC (and ECL) relative to the working concentration from **Table S1**. For example, 40% EggPC means that the Lipid-IPA mix contains 0.52 mM of EggPC phospholipids (40% of 1.3 mM) and 1.8 mM of ECL phospholipids (60% of 3 mM).

To determine whether IIMPs specific to different lipid kingdoms can coexist in hybrid membranes, such as the one that could be developed in engineered biochemical systems^19^, we devised hybrid SLBs made of EggPC and ECL. We measured the shift in frequency Δf of blank TXTL reactions (plasmid P70a-*T7rnap* only) and of TXTL reactions expressing *ah-egfp* and *mscL* (**Figure 3d, Figure S12**). The shift in frequency Δf was measured after eight hours of incubation, taken as an amount of time sufficient for the TXTL reaction and interaction with the SLB to reach equilibrium. We first established that at any ratio of phospholipids, the hybrid membrane formed remains devoid of non-specific adsorption. In contact with the blank TXTL reaction, at any EggPC ratio, the frequency shift stabilized within 2 hours of incubation (**Figure S12a**). We also observed a slight linear increase in the Δf with the increase of the EggPC ratio, which accounts for slight nonspecific interactions of the TXTL reaction with *E. coli* SLBs. For AH-eGFP, we observed a sigmoidal decrease of Δf as the EggPC ratio increased, whereas for MscL Δf increased linearly with the increase of the EggPC ratio, consistent with the respective specificity of AH-eGFP and MscL integration into EggPC and ECL. Considering the cooperativity of the assembly of multimeric complexes in the plasma membrane of bacteria^42^, the sigmoidal dependence for AH-eGFP is expected^43^, while the linear dependence observed for MscL is surprising. When the MscL TXTL reaction was incubated onto DOPE-DOPC hybrid SLBs, MscL exhibited a sigmoidal increase with the increase of the DOPC ratio (**Figure S13**). These experiments underline the existence of different self-assembly and integration mechanisms, depending on the presence of lipids, such as DOPG or CL, present in ECL but absent in DOPE^44–46^.

### SLBs’ lipid compositions can be broadly tuned

Although PC (eukaryotes) and PE (bacteria) are usually the predominant phospholipids in natural cell membranes^47–49^, other headgroups are similarly essential for the function of IIMPs^27^. For example, some IIMPs only anchor to specific phospholipid heads other than PE or PC^27^, while other IIMPs require the presence of anionic phospholipids to bind^50^. While making SLBs from a mixture of different phospholipids is technically feasible with the SALB method (**Figure 4a**), one must determine the degree of nonspecific adsorption of TXTL on the SLBs. We measured the concentrations of DOPG, CL, and DPPS (1,2-dipalmitoylphospho-L-serine) into three of our base SLBs (DOPC, EggPC, and DOPE) for which nonspecific TXTL adsorption is absent (**Figure 4a**). We used the DPPS sensing protein LactC2 to show that one can adjust quantitatively the amount of an added lipid to an SLB.

**Figure 4.**
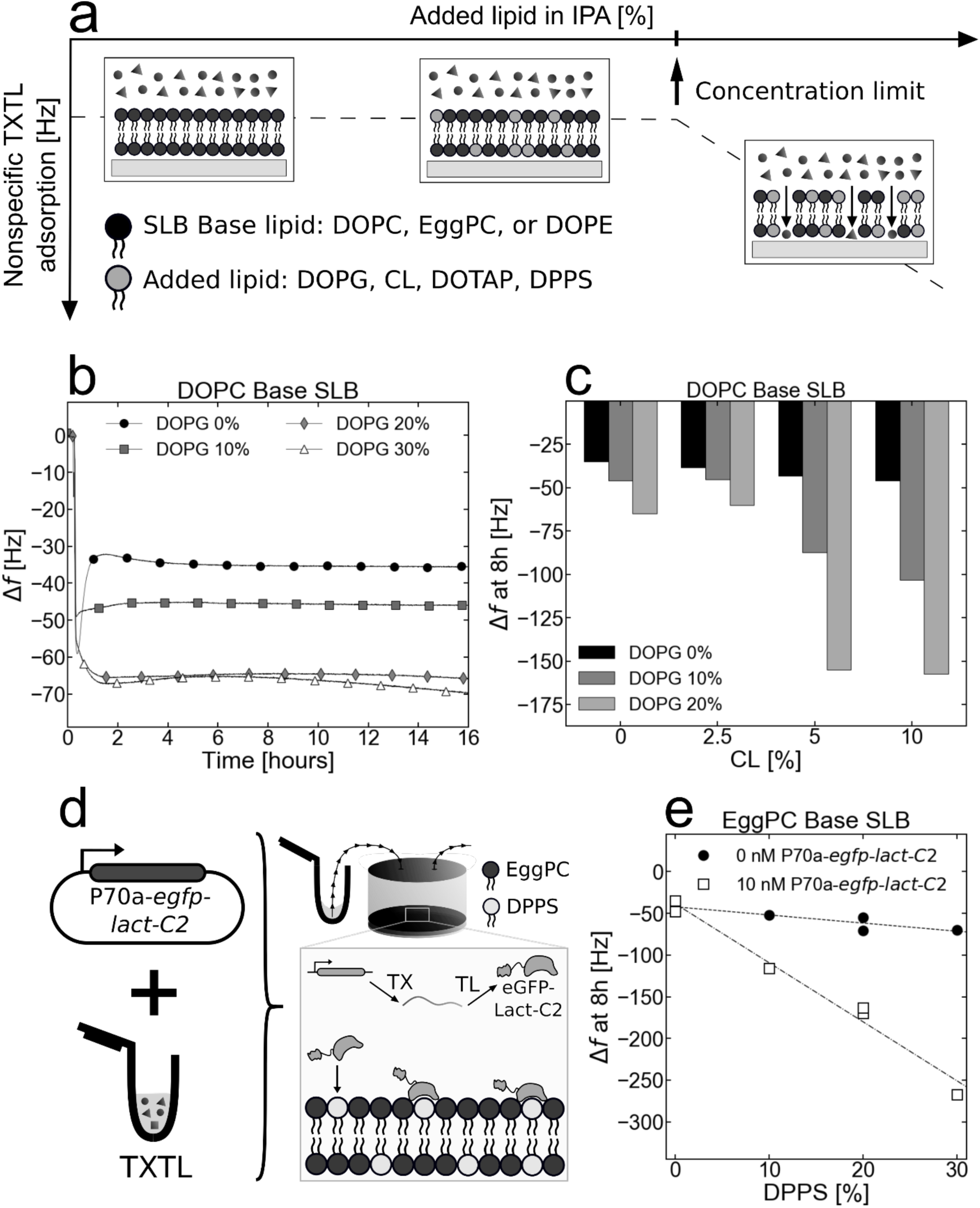
Phospholipids that cannot form stable SLBs in normal TXTL conditions can be added to ‘base lipid’ SLBs such as DOPC, EggPC, and DOPE SLBs. **a.** Graphical illustration of the ‘added lipid’ concentration limit. Although phospholipids with various headgroups can be added to base SLBs, there is an observed limit to the added lipid concentration in the phospholipid-IPA mix before the SLB-sensor system incurs nonspecific adsorption from a blank TXTL reaction. **b.** Adsorption kinetics of a blank TXTL reaction (P70a-*T7rnap*, 0.15 nM) in contact with DOPC/DOPG SLBs of different ratios. **c.** The frequency change after 8 hours of a blank TXTL reaction (P70a-*T7rnap*, 0.15 nM) incubating over a DOPC Base SLB as a function of added CL and DOPG. **d.** Cell-free synthesis of eGFP-Lact-C2 (plasmid P70a-*egfp-lact-C2*, 5 nM) into the QCMD module. The *in situ* synthesized eGFP-Lact-C2 interacts with the DPPS phospholipids. **e.** The frequency change after 8 h of either a blank (no DNA added) or eGFP-Lact-C2 (P70a-*egfp-lact-C2*, 5 nM) TXTL reaction being in contact with EggPC/DPPS SLBs fabricated with varying SALB DPPS concentrations.

The added lipid percentage of the lipid-IPA mixture is defined as the ratio of the molarity of the added lipid over that of the base lipid. Using a blank TXTL reaction (plasmid P70a-*T7rnap*, 0.15 nM), we found that adding DOPG up to 30% to either DOPC or DOPE SLBs did not produce nonspecific TXTL adsorption (frequency drifts of less than 1 Hz/hour) (**Figure 4b and Figure S14**). We also found that up to 10% of CL in DOPE SLBs (**Figure S15**) and in DOPC SLBs (**Figure S16a**) did not produce non-specific adsorption. The conditions for which nonspecific TXTL adsorption is not observed when both DOPG and CL are added to a DOPC SLB were also determined (**Figure 4c, Figure S16b, c**). As the concentration of CL in the lipid-IPA mix is increased, the amount of DOPG must be decreased to prevent nonspecific TXTL adsorption. The limit concentration of each of the tested added lipids was determined by following the same procedure (**Table S1**).

LactC2 is a mammalian signaling protein that binds specifically to PS phospholipids^23,51,52^. When eGFP-LactC2 was synthesized (plasmid P70a-*egfp-lactc2*, 5 nM) on DPPS-EggPC SLBs, the shift in frequency Δf was linearly proportional to the amount of DPPS added to the IPA-EggPC-DPPS solution used to form the SLB (**Figure 4d**, **e, Figure S17**). Because more eGFP-LactC2 proteins were synthesized than DPPS sites in the SLB available for binding **(Figure S17b)**, this experiment shows that the amount of DPPS in an EggPC base SLB corresponds to the proportion of DPPS used to make the SLB.

### The *E. coli* Min system produces oscillations on SLBs

To determine whether the TXTL-QCMD approach enables the sensing of membrane-based dynamical patterns on and greater than mesoscopic scales, we assayed the *E. coli* Min system that uses the inner membrane to position the division machinery via pole-to-pole dynamical oscillations^53^. *In vitro*, the three Min proteins MinD, MinE, and MinC create a myriad of oscillatory patterns in vesicles^54–57^ and on SLBs^58–61^. These patterns are the result of the reaction-diffusion dynamics of MinD and MinE. Upon ATP binding, MinD forms a dimer that binds to the inner bacterial membrane, which accelerates MinD recruitment at the membrane. MinE, which forms a dimer in the cytoplasm, binds to the MinD dimer at the membrane and catalyzes the hydrolysis of the ATP bound to MinD, effectively releasing MinD from the membrane^62–65^. While not essential for oscillations^54^, MinC is known to bind to MinD and ‘ride’ the MinD waves^66^. Visualizing MinCDE dynamical patterns requires fluorescent tagging of the proteins, which can interfere with the function of proteins^67^.

First, we performed a set of experiments in which we only synthesized MinD onto the SLBs (10 nM linear P70a-*minD*). Previous studies have shown that anionic phospholipids, such as PG and CL, facilitate MinD interaction with the membrane^68^. We added either DOPG or CL to a DOPC SLB (**Figure 5a**). No oscillations were observed on pure DOPC SLBs, while on the DOPG-DOPC and CL-DOPC SLBs the synthesized MinD induced oscillations with 5.5 min and 6.7 min mean periods respectively in the first two hours (**Figure 5b**). While MinC is not necessary for the Min oscillations^54^, it has been reported that both MinD and MinE are required to form spatio-temporal patterns^3^. Proteomics of our TXTL system showed trace amounts of MinD, whereas MinE and MinC were not detected^35^. Blank TXTL reactions never produced spontaneous oscillations on SLBs. The synthesized MinD oscillations exhibited two different modes: an unstable and large-amplitude mode in the first 2 hours of incubation and a stable small-amplitude mode for the rest of the reaction (**Figure S18a, b, c**). In both modes, the period was around 10 minutes, longer than the 1 minute periods observed in confined systems such as *E. coli* bacteria^69^ or micron-sized microwells^70^. The oscillations stopped with an increase in Δf of 2-10 Hz depending on the replicate, which we interpret as the final unbinding of MinD from the SLB after the ATP that fueled the oscillations has been depleted (**Figure S18d**).

**Figure 5.**
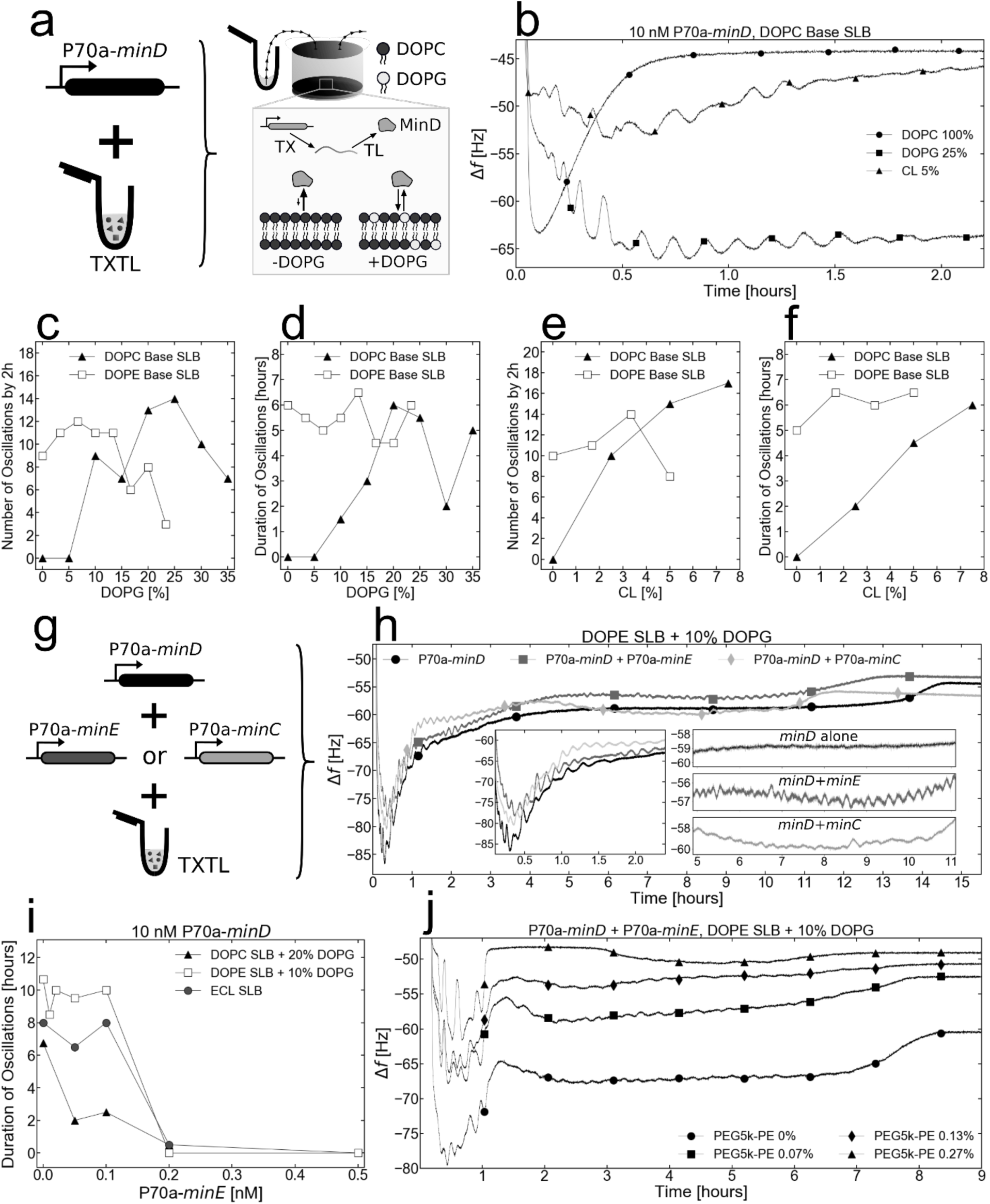
*In situ* TXTL synthesis of Min proteins induces coherent long-range mass oscillations across the SLB-Sensor system. **a.** Cell-free synthesis of MinD (linear P70a-*minD*) into the QCMD modules on top of SLBs. The cartoon shows a very simplified model of the synthesized MinD binding kinetics when the reaction is in contact with either a pure DOPC SLB or a DOPC/DOPG SLB. **b.** Adsorption kinetics of MinD with either a pure DOPC SLB, a DOPG/DOPC SLB, or a CL/DOPC SLB. The curves are labeled with a few symbols, the lines are experimental points. **c.** Number of oscillation peaks within the first 2 hours as a function of the relative DOPG concentration during SALB for a DOPE and DOPC Base SLB. **d.** Duration of oscillations as a function of the relative DOPG concentration during SALB for a DOPE and DOPC Base SLB. **e.** Number of oscillation peaks within the first 2 hours as a function of the relative CL concentration during SALB for a DOPE and DOPC Base SLB. **f.** Duration of oscillations as a function of the relative CL concentration during SALB for a DOPE and DOPC Base SLB. **g.** Cell-free synthesis of MinD with either MinE (linear P70a-*minE*) or MinC (linear P70a-*minC*) into the QCMD modules on top of SLBs. **h.** Adsorption kinetics of MinD (P70a-*minD*, 10 nM), MinDE (P70a-*minD*, 10 nM and P70a-*minE*, 0.1 nM), and MinCD (P70a-*minD*, 10 nM and P70a-*minC*, 5 nM), on a 10% mol. ratio DOPG/DOPE SLB. The left inlet shows a close-up of the oscillations during the first 2.4 hours. The right inlet shows a close-up of the oscillations between t =5 hours and t = 11 hours. The curves are labeled with a few symbols, the lines are experimental points. **i.** Duration of oscillations as a function of P70a-*minE* concentration in the TXTL reaction for a DOPG/DOPE SLB, a DOPG/DOPC SLB, and an *E. coli* lipids (ECL) SLBs. **j.** Adsorption kinetics of a MinDE (P70a-*minD*, 10 nM and P70a-*minE*, 0.1 nM) on a 10% mol. ratio DOPG/DOPE SLB with a range of PEG5k-PE added during SALB. The curves are labeled with a few symbols, the lines are experimental points.

Next, we determined the ranges of concentrations of DOPG and CL that can be added to either a pure DOPC (not found in *E. coli* membranes) or pure DOPE (major phospholipid in *E. coli* membranes) SLB for which the Min oscillations were observed when MinD (10 nM linear P70a-*minD*) only was synthesized in the TXTL reaction. To track the effect of DOPG concentrations, we counted the number of oscillations during the first mode (larger oscillations), and the duration of the oscillations during the second mode (smaller oscillations). In this series of experiments, the behavior of the initial mode was strongly dependent on the charge of the SLB, with both the DOPE and DOPC base SLBs exhibiting optimal DOPG molar ratios of 5-10% and 20-25% respectively (**Figure 5c, Figure S19, S20**). The difference in the optimal DOPG percentage can be partially attributed to the difference in the availability of the positive charge in the zwitterionic headgroups of DOPE and DOPC. When the reaction duration was measured, we noticed that the DOPE SLBs were significantly more robust to variations in the DOPG concentration compared to the DOPC SLB (**Figure 5d, Figure S21, S22**). Similarly, as we increased the concentration of CL in the SLB, we observed an increase of the number of oscillations in the first 2 hours alongside the increase of oscillation lifetime for the DOPC SLB cases (**Figure 5e, f**). Changes in the CL concentration in DOPE base SLB did not significantly affect the behavior of the pattern-forming reactions. Overall, the DOPE base SLB provided more robustness to lipid composition changes compared to the DOPC base SLBs for observing oscillations. When DOTAP, an artificial cationic phospholipid, was added to DOPC, DOPG-DOPC, DOPE, and DOPG-DOPE SLBs, the oscillations became weaker as the concentration of DOTAP increased (**Figure S23, S24, S25**). This shows that Min patterns are sensitive to the charges present in the SLB, with negative charges helping their formation, while positive charges inhibit oscillations. This result can be considered together with the fact that the *E. coli* inner membrane has a highly nonuniform density of PG and CL, with larger densities localized at the poles, especially for CL^71–73^. Min patterns in *E. coli* must be able to maintain oscillatory order in these locally different phospholipid compositions simultaneously, thus requiring the patterns to be robust to anionic phospholipid variation.

We co-synthesized either MinC or MinE with MinD in the presence of a DOPE-based SLB with 10% DOPG (**Figure 5g**) to determine the effect of each on the MinD oscillations. In the presence of MinC (2 nM linear P70a-*minC*), the period of the oscillations remained unchanged while their amplitude increased. Adding MinE in some of the reactions produced a doubling of the mass oscillation period **(Figure 5g)**, which is likely an oscillatory regime in which MinE is recruited by MinD on some regular distance as observed in previous MinDE SLB experiments^59^. This regime did not occur in all the MinDE experiments that we tested but revealed patterns that can exist on the sensor when both *minD* and *minE* were expressed at optimal concentrations. To determine the concentrations of P70a-*minE* for which the pattern-forming reaction is the longest, we performed a series of experiments in which the *minD* expressing DNA concentration was fixed (10 nM linear P70a-*minD*), while the concentration of the *minE* expressing DNA was varied (linear P70a-*minE*). These *minE* DNA ranges were performed for the optimal DOPG concentrations of both the DOPE (10% DOPG) **(Figure S26)** and DOPC (20% DOPG) **(Figure S27)** base SLBs and for the pure ECL SLB **(Figure S28)**. The DOPG/DOPC SLB demonstrated a decay in the reaction time as the MinE-expressing DNA was increased in concentration. DOPG/DOPE and ECL SLBs demonstrated robustness to the MinE variation up to 0.1 nM of added P70a-*minE* and no prolonged oscillations with P70a-*minE* concentrations greater than 0.1 nM (**Figure 5i**). These results support that PE phospholipids play an integral role in the robustness of Min patterns, and that PG is not the only critical headgroup for Min oscillations.

Natural membranes are crowded with many membrane proteins. To address the effect that molecular crowding has on the Min patterns, we added PEG5000-PE (PE phospholipid with a PEG moiety of molecular mass 5000 g/mol) at various concentrations into a DOPE SLB with 10% DOPG. We expressed both *minD* (10 nM linear P70a-*minD*) and *minE* (0.1 nM linear P70a-*minE*) at optimal concentrations in the QCMD module. As we increased the amount of PEG5000-PE in the SLB, we observed an increase in the amplitude of the early oscillations (**Figure 5j**). Membrane molecular crowding accelerates the binding of MinD to the SLB, which can be reasoned by the fact molecular crowding facilitates the formation of MinD multimers^74–77^. Molecular crowding decreased the total reaction duration and in the 0.27% PEG5000-PE case caused a two-hour long transient disappearance of oscillations after the large amplitude oscillations in the first hour. It is important to note that producing mass oscillations across the entire sensor requires that the MinD proteins collectively bind and unbind from the SLB in-phase and the transient disappearance of the oscillations might indicate that the Min patterns are exhibiting energetically costly chaotic dynamics^61,78,79^ during which the MinDE system is assessing a more ordered state^80,81^.

The *in situ* synthesized MinCDE proteins produce mass oscillations of proteins adsorbed onto the SLB that can be studied as a function of the SLB composition. In some cases, MinDE oscillations on the 1.2-cm QCMD sensor were observed for more than 12 hours. We found that, while anionic phospholipids such as DOPG and CL are necessary to observe MinD spatio-temporal patterns in DOPC base SLBs, those patterns can be observed in pure DOPE SLBs. Although the literature suggests that MinE is critical for Min oscillations, we found that MinE is not necessary for those patterns to form, to our surprise. The data suggest that the concentric circular Min waves on the circular sensor have a wavelength of about 100 µm, which is on the order of the wavelength measured previously in a similar planar configuration^58^ **(Figure S18e)**.

### Type II Zorya antiphage defense system is membrane-associated

Recently, a plethora of novel microbial defense systems have been discovered in genomic defense islands^24,82^ and in phages^83^. Yet, their mechanisms of immunity largely remain unknown. One of such defense systems, type II Zorya, consists of ZorA, ZorB, and ZorE proteins that together protect their host *E. coli* from phage infection via an unknown mechanism^24^. ZorA and ZorB are likely IIMPs, as they contain predicted membrane interacting domains **(Figure 6a)** and are homologous to the flagellar motor proteins MotA and MotB^24^. Here, we demonstrated that the native Zorya system protects *E. coli* ATCC 8739 against phages T7 and PhiX174 (**Figure S29a**). However, tagging the individual Zorya proteins for determining their localization and for subsequent purification disrupted their protective function (**Figure S29b, c, d**), making this novel defense system particularly challenging to study. To overcome these limitations, we used the TXTL/QCMD approach to unravel the interactions of the Zorya proteins with lipid bilayers and to shed light on its mechanism of immunity.

**Figure 6.**
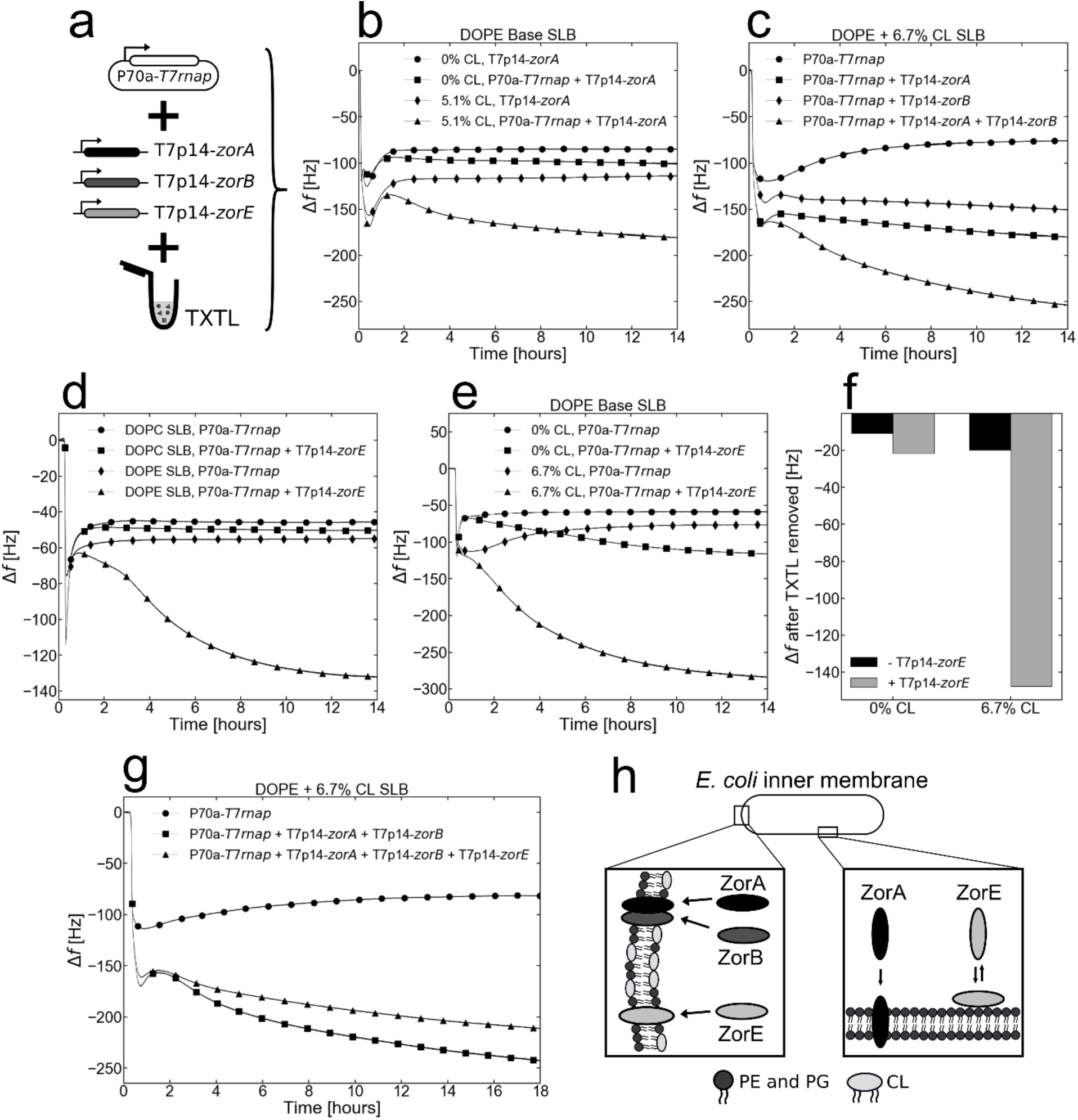
Type II Zorya defense proteins interact with cardiolipin-rich lipid membranes. **a.** ZorA, ZorB, and ZorE were synthesized via the T7 transcriptional activation cascade (plasmid P70a-*T7rnap*, 0.15 nM, linear T7p14-*zorA*, T7p14-*zorB*, T7p14-*zorC*, 10 nM). The TXTL reactions were immediately pushed into the QCMD modules with SLBs on the sensor. **b.** Adsorption kinetics of the blank (P70a-*T7rnap*, 0.15 nM) and ZorA (P70a-*T7rnap*, 0.15 nM and T7p14-*zorA*, 10 nM) conditions with either a pure DOPE or DOPE + 5.1% CL SLBs. **c.** The adsorption kinetics of the blank (P70a-*T7rnap*, 0.15 nM), ZorA alone (P70a-*T7rnap*, 0.15 nM and T7p14-*zorA*, 10 nM), ZorB alone (P70a-*T7rnap*, 0.15 nM and T7p14-*zorB*, 10 nM), and ZorA and ZorB together (P70a-*T7rnap*, 0.15 nM, T7p14-*zorA*, 10 nM and T7p14-*zorB*, 10 nM) with a DOPE + 6.7% CL SLB. **d.** The adsorption kinetics of the blank (P70a-*T7rnap*, 0.15 nM) and ZorE (P70a-*T7rnap*, 0.15 nM and T7p14-*zorE*, 10 nM) conditions with either a pure DOPC or pure DOPE SLBs. **e.** The adsorption kinetics of the blank (P70a-*T7rnap*, 0.15 nM) and ZorE (P70a-*T7rnap*, 0.15 nM and T7p14-*zorE*, 10 nM) conditions with either a pure DOPE or DOPE + 6.7% CL SLBs. **f.** The remaining change in frequency in the conditions from **e.** after the SLB with the interacting proteins are flushed with Tris NaCl for 1 hour. **g.** The adsorption kinetics of the blank (P70a-*T7rnap*, 0.15 nM), ZorAB (P70a-*T7rnap*, 0.15 nM, T7p14-*zorA*, 10 nM, and T7p14-*zorB*, 10 nM), and ZorABE (P70a-*T7rnap*, 0.15 nM, T7p14-*zorA*, 10 nM, T7p14-*zorB*, 10 nM, and T7p14-*zorE*, 5 nM) conditions with a DOPE + 6.7% CL SLB. **h.** An illustration of the hypothetical localization of the Zorya proteins in the host *E. coli* inner membrane. All Zorya proteins preferentially interact with the CL-rich polar regions of the inner membrane. In addition, ZorA likely has a weak interaction with the equatorial region of the inner membrane and ZorE likely has a reversible interaction with the equatorial region of the inner membrane.

ZorA, ZorB, and ZorE were synthesized through the T7 transcriptional activation cascade **(Figure 6a)**. The interaction of ZorA with SLBs was enhanced when CL was added to DOPE SLBs (**Figure 6b**). However, adding DOPG to DOPE SLBs did not result in improved ZorA interaction (**Figure S30**). Expectedly, ZorA did not interact with a DOPC SLB suggesting a preference for bacterial lipids (**Figure S31**). When ZorA and ZorB were synthesized together, they produced a larger QCMD frequency shift than either of them alone, which indicates that they form a complex at the membrane (**Figure 6c**). On an ECL SLB, ZorA produces a small frequency shift with or without ZorE (**Figure S32**). The kinetics of such shift is fast, indicating that the concentration of CL in ECL is small compared to the one tested **(Figure 6 b, c, e)**.

Although ZorE is not predicted to have membrane interacting domains^24^, ZorE protein interacted with a pure DOPE SLB but not with a pure DOPC SLB (**Figure 6d**). As with ZorAB, ZorE interacted more strongly with a DOPE/CL SLB than a pure DOPE SLB (**Figure 6e**). To determine the strength of the interaction with the SLBs, we flushed Tris-NaCl though the QCMD chamber above the SLBs at the end of the TXTL reaction. We observed that the ZorE frequency change was preserved only when CL was present in the SLB. In the condition with a pure DOPE SLB, it appears that all the synthesized ZorE proteins interacting with the SLB have been removed during flush (**Figure S33**, **Figure 6f**). This suggests that ZorE interacts weakly on the surface with a pure DOPE membrane but interacts irreversibly with membranes containing cardiolipin. Finally, the co-synthesis of ZorE with ZorAB did not result in a stronger interaction with a DOPE/CL SLB than ZorAB alone (**Figure 6g**).

Overall, our results suggest that the ZorABE proteins likely localize at the poles of *E. coli* where cardiolipin is most dense (**Figure 6h**). ZorE likely diffuses along the surface of the inner membrane and embeds into the membrane upon finding cardiolipin-rich domains. Considering that Zorya protects *E. coli* from T7 and that T7 preferably binds *E. coli* cells at the poles^84^, it is possible that the ZorA and ZorB proteins disrupt the membrane potential that T7 uses to inject the first segment of its genome into the cell^85^. Alternatively, ZorAB may sense membrane depolarization caused by T7 infection and activate ZorE, which contains a predicted HNH domain **(Figure 6a)**, as a putative downstream effector to enact immunity.

## Conclusion

The QCMD-TXTL approach for characterizing IIMPs’ interactions with lipid bilayers has several advantages compared to the current methods **(Table 1)**. It is fast, non-disruptive, and has an excellent degree of reproducibility. It enables assaying single or concurrently *in situ* synthesized IIMPs on membranes with arbitrarily complex lipid compositions on physical scales that tolerate the formation of large dynamical patterns. The QCMD-TXTL method proves to be useful for unraveling basic underpinning details about how IIMPs interact with different lipids and assemble at or into membranes. This method also provides information for engineering synthetic membranes capable of hosting IIMPs that naturally reside in different lipid environments. Taken altogether, these features make the QCMD-TXTL approach a highly favorable method for characterizing IIMPs-membranes interactions. We anticipate that it could be useful for the bottom-up engineering of systems such as synthetic cells^19,86–89^.

**Table 1.**
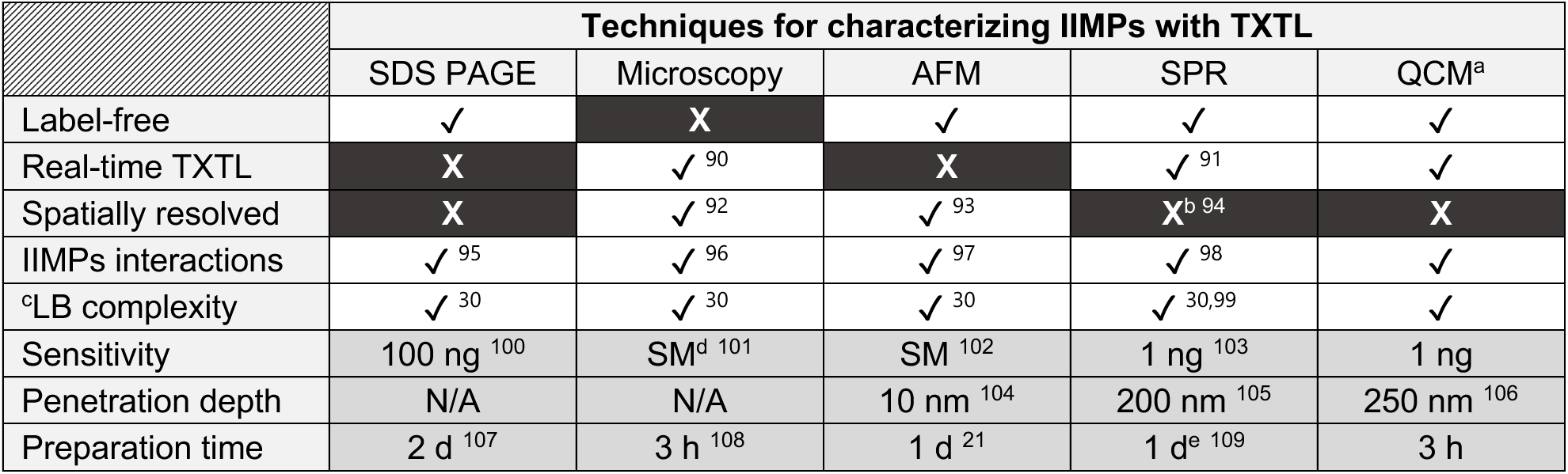
Comparison of techniques for cell-free synthesized IIMP characterization. ^a^: the other information for QCMD is from this study. ^b^: SPR can have spatial resolution, but this modality is not used for IIMP characterization. ^c^: the lipid bilayer (LB) complexity refers to whether any type of lipid compositions can be achieved. ^d^: single molecule (SM). ^e^: Potentially, the preparation time of SPR can be reduced to the same time as QCM experiments, but no studies utilizing the SALB-TXTL approach with SPR have been reported.

In the field of *E. coli* cell-free expression systems, it is established that *E. coli* inner membrane proteins integrate into lipid membranes without secretion mechanisms via a process coupled to translation^110,111^. This assumption is supported by the fact that IIMPs also integrate into lipid membranes when they are synthesized with the PURE system deprived of any secretion components^112^. In lysate-based TXTL systems, it is not clear whether soluble parts of the secretion mechanism present in the lysate (*e.g.*, SecA is present in *E. coli* lysates^35^) help IIMPs integrate into membranes. In this work, we used an *E. coli* TXTL system, commercially available under the name of myTXTL (Arbor Biosciences). Other TXTL systems could be used if protein synthesis yields are large enough, which should be on the order of at least 100 nM based on our measurements and estimations **(Figure S11)**. We anticipate that most of the IIMPs from bacteria could be assayed using TXTL systems from *E. coli* or other bacteria, many of which have been reported in the literature^113^. Studying IIMPs from higher organisms with the QCMD-TXTL method remains to be investigated and is likely to be dependent on the complexity of the post-translation modifications of each IIMP. The scope of the IIMPs assayed in this work was chosen to demonstrate (i) that the QCMD-TXTL method corroborates many observations previously reported, (ii) and that novel insights and discoveries into the IIMPs-membranes’ interactions can be rapidly obtained. For instance, deciphering the Zorya system supports that many other novel defense systems could be deciphered using this approach.

The QCMD provides two signals, the change in frequency and the dissipation. In our work, we only used the change in frequency change Δf as it is proportional to the mass added to the SLB. The dissipation term gives information about the viscoelastic changes of the lipid bilayer. As opposed to the frequency change, however, the dissipation term is much more difficult to understand and has not been subject to a comprehensive characterization in the literature. Extensive studies based on model systems would be necessary to make dissipation a truly interpretable term.

We focused our work on using planar SLBs formed on the QCMD sensor. We showed that this configuration enables probing IIMPs’s interactions with membranes over a broad variety of lipids, including lipids localized *in vivo* like cardiolipin. This configuration does not enable looking at specific variables found in living systems, such as membrane curvature for example. Instead of making flat SLBs, vesicles could be attached to the sensor, which has been done previously without TXTL^114^. We anticipate that dynamical patterns, such as the Min system, would be harder to achieve in this configuration, which would also require passivating the sensor to prevent nonspecific adsorption. Alternatively, SLBs can be fabricated on top of silica nanoparticles of controlled radii to mimic membrane curvature^115^.

## Methods

### Reagents

All the phospholipids were obtained from Avanti Polar Lipids in powder form. The phospholipids were dissolved in IPA (isopropyl alcohol, ThermoFisher Scientific, A416S-4) at the following stock concentrations: EggPC – 100 mg/mL (840051P), DOPC (1,2-dioleoyl-sn-glycero-3-phosphocholine) – 100 mg/mL (850375P), ECL (100500P) – 50 mg/mL, DOPE (1,2-dioleoyl-sn-glycero-3-phosphoethanolamine) – 25 mg/mL (850725P), DOPG (1,2-dioleoyl-sn-glycero-3-phospho-(1’-rac-glycerol)) – 10 mg/mL (840475P), CL (1’,3’-bis[1,2-distearoyl-sn-glycero-3-phospho]-glycerol) – 5 mg/mL (710334P), DOTAP (,2-dioleoyl-3-trimethylammonium-propane) – 10 mg/mL (890890P), DPPS (1,2-distearoyl-sn-glycero-3-phospho-L-serine) - 50 mg/mL (840029P), PEG5000-PE (1,2-dipalmitoyl-sn-glycero-3-phosphoethanolamine-N-[methoxy(polyethylene glycol)-5000]) – 0.5 mg/mL (880200P). To prepare the working IPA-lipid mixes, an appropriate volume of the stock phospholipids was diluted in IPA to the desired concentration, vortexed at medium-high speed for ten seconds, and immediately used for SALB.

### Cell-free transcription-translation

Cell-free gene expression was carried out using an *E. coli* TXTL system described previously^26^, with one modification. We used the strain BL21-Δ*recBCD* Rosetta2 in which the *recBCD* gene set is knocked out to prevent the degradation of linear DNA^116^. The preparation and usage of the TXTL system were the same as reported before^26,117^. Briefly, *E. coli* cells were grown in a 2xYT medium supplemented with phosphates. Cells were pelleted, washed, and lysed with a cell press. After centrifugation, the supernatant was recovered and preincubated at 37 °C for 80 min. After a second centrifugation step, the supernatant was dialyzed for 3 h at 4 °C. After a final spin-down, the supernatant was aliquoted and stored at −80 °C. The TXTL reactions comprised the cell lysate, the energy and amino acid mixtures, maltodextrin (30 mM) and ribose (30 mM), magnesium (2-5 mM) and potassium (50-100 mM), PEG8000 (1-2 wt%), water and the DNA to be expressed. The TXTL reactions were incubated at 30 °C in QCMD chambers (40 µl reactions). The TXTL reactions were incubated at 30 °C on 96 well plates (2 µl reactions) for the measurement of the kinetics of deGFP synthesis.

### Reporter protein quantification

AH-eGFP and LactC2-eGFP were quantified on a Biotek H1M plate reader using a calibration curve. The calibration curve was determined using pure eGFP (Cell Biolabs, STA-201) following a procedure described earlier^26^.

### DNA preparation

All the DNA sequences are reported in the supplementary DNA file. We either used plasmids or linear DNAs. DNAs were obtained either by PCR or purchased (Twist Biosciences, IDT). DNA stock solutions were quantified with a spectrophotometer (Thermofisher Scientific, NanoDrop 2000). Genes were expressed either from the strong *E. coli* promoter P70a or the T7 transcriptional activation cascade^17,25,26^.

### QCMD sensors and modules preparation

A QSense Analyzer (Biolin Scientific, Gothenburg, Sweden) was used for the experiments. The QSense Analyzer has four channels that can be used independently and concurrently. The preparation of the instrument for an experiment consisted of a 400 µl/min flush of at least 20 mL of a 1% wt SDS solution, then 20 mL of autoclaved deionized water, then air until the tubing was empty. After the four QCMD modules were dried, they were disconnected from the QSense Analyzer mount and opened. After removing the sensors (Biolin Scientific, QSX303), the QCMD modules were dried with nitrogen. The sensors were dried with nitrogen as well and plasma cleaned (Harrick Plasma, PDG-32) at low RF power for 10 minutes. The plasma-cleaned sensors were then reinserted into the dry QCMD modules. The QCMD modules were closed and connected to the QSense Analyzer mount. Using the QSoft software, the bare sensors were calibrated and the QCMD modules were ready.

### QCMD SLB preparation and TXTL reaction

The QCMD modules were maintained at 30 °C throughout the whole experiment. First, a Tris NaCl Buffer (10 mM Tris, 150 mM NaCl, 7.5 pH) was flushed at 100 µl/min through all the modules until the resonance frequency of the sensors stabilized, which takes 40-60 minutes. Next, IPA was flushed until signal stabilization, which takes around 20 minutes and produces a change in frequency of approximately -75 to -80 Hz. We then flushed the IPA-lipid mixes at 100 µl/min for 20 minutes. After stabilization, the signal varies by -5 to -10 Hz, depending on the lipids used, usually within 5-10 minutes, except for ECL which requires the full duration of the flush. Next, we flushed the Tris NaCl buffer at 100 µl/min until signal stabilization to complete the formation of the SLB. This step takes up to 40 minutes and produces a change of frequency of the order of 30 to 50 Hz, depending on the lipids used. Finally, we pushed the TXTL reactions (up to four different conditions, one per module) at 25 µl/min. The pumps were stopped and the TXTL reactions were incubated in contact with the SLB-sensor system for 3 - 20 h.

### QCMD data analysis

The analysis of QSense Analyzer data was performed on the 7th overtone of the resonance of the sensor due to its lowest sensitivity to variations in the mounting of the sensor, following the manufacturer’s recommendation. The frequency at the end of the second Tris NaCl flush during the SLB preparation was reset as Δf = 0. The frequency output of the seventh harmonic was divided by 7 to reflect the effect the TXTL reactions had on the fundamental frequency changes of the sensor.

### Zorya knockout mutants

Zorya genes *zorABE* were deleted from the genome of *Escherichia coli* DSM 1576 (ATCC 8739) using lambda red recombination as described^118^. The genes were deleted in the region of the chromosome between 4,265,744 and 4,269,528 bp (NCBI Ref. CP000946).

### Plaque assay

Double layer plaque assay was used to determine phage titers. *E. coli* were cultivated in LB medium until the OD_600_ of 0.55 before centrifugation. The pellet was resuspended in cold LB, mixed with soft agar, and the appropriate phage dilution. This mixture was added to an agar plate and solidified. Subsequently, phage dilutions were directly spotted onto the plate. The plates were incubated at 37°C until visible plaques developed within 4 to 18 hours.

## Associated Content

Supplementary Figures S1-S33, Table S1.

Supplementary File Table S2.

## Author Information

## Corresponding Author

Vincent Noireaux - School of Physics and Astronomy, University of Minnesota, Minneapolis, MN 55455, USA https://orcid.org/0000-0002-5213-273X

## Authors

Aset Khakimzhan – School of Physics and Astronomy, University of Minnesota, Minneapolis, MN 55455, USA

Ziane Izri – School of Physics and Astronomy, University of Minnesota, Minneapolis, MN 55455, USA

Seth Thompson – School of Physics and Astronomy, University of Minnesota, Minneapolis, MN 55455, USA

Oleg Dmytrenko – Helmholtz Institute for RNA-based Infection Research (HIRI), Helmholtz-Centre for Infection Research (HZI), 97080 Würzburg, Germany

Patrick Fischer – Helmholtz Institute for RNA-based Infection Research (HIRI), Helmholtz-Centre for Infection Research (HZI), 97080 Würzburg, Germany

Chase Beisel – Helmholtz Institute for RNA-based Infection Research (HIRI), Helmholtz-Centre for Infection Research (HZI), 97080 Würzburg, Germany; Medical Faculty, University of Würzburg, 97080 Würzburg, Germany

## Author contributions

A.K., Z. I., S. T., O.D., performed the experiments. A.K., Z. I., S. T., O.D., C.L.B., and V.N. designed the experiments, analyzed the data, and wrote the manuscript. V.N. edited the manuscript.

## Acknowledgments

The authors thank David Garenne for his help in the preparation of the TXTL system used in this work. This work and the materials are based on funding provided by the National Science Foundation (BBSRC-NSF/BIO 2017932 to V.N.) and the Deutsche Forschungsgemeinschaft SPP 2330 program (BE 6703/2-1 to C.L.B.).

