## Supplementary Figures S1-S33, Table S1 for "Cell-free expression with a quartz crystal microbalance enables rapid, dynamic, and label-free characterization of membrane-interacting proteins"


| Item | Title | Pages |
|---|---|---|
| Figure S1 | TXTL lysate does not contain living E. coli cells | 2 |
| Figure S2 | Synthesis of deGFP from linear templates with a P70a promoter or a T7 cascade | 3 |
| Figure S3 | SALB procedure with and without lipids during step III | 4 |
| Figure S4 | Nonspecific TXTL adsorption for model SLBs | 5 |
| Figure S5A | MscL TXTL adsorption replicates with ECL SLB | 6 |
| Figure S6 | AH-eGFP TXTL adsorption replicates with EggPC SLB | 7 |
| Figure S7 | Specific adsorption of MscL and AH-eGFP into DOPC and DOPE SLBs | 8 |
| Figure S8 | Pre-synthesized MscL does not integrate into SLBs | 9 |
| Figure S9 | Pre-synthesized AH-eGFP does integrate into SLBs | 10 |
| Figure S10 | Post-QCMD fluorescence measurements | 11 |
| Figure S11 | Mass sensitivity of the QCMD | 12 |
| Figure S12 | Adsorption of blank, AH-eGFP, and MscL TXTL reactions onto ECL-EggPC SLBs | 13 |
| Figure S13 | Adsorption of blank and MscL TXTL reactions into DOPE-DOPC SLBs | 14 |
| Figure S14 | Nonspecific TXTL adsorption into DOPG-DOPE SLBs | 15 |
| Figure S15 | Nonspecific TXTL adsorption into CL-DOPG-DOPC SLBs | 16 |
| Figure S16 | Nonspecific TXTL adsorption into CL-DOPE SLBs | 17 |
| Figure S17 | eGFP-LactC2 adsorption kinetics and approximate endpoint concentrations | 18 |
| Figure S18 | Modes and wavelengths of oscillations produced by MinD | 19 |
| Figure S19 | MinD adsorption kinetics in DOPG-DOPC SLBs | 20 |
| Figure S20 | MinD adsorption kinetics in DOPG-DOPE SLBs | 21 |
| Figure S21 | MinD adsorption kinetics in CL-DOPC SLBs | 22 |
| Figure S22 | MinD adsorption kinetics in CL-DOPE SLBs | 23 |
| Figure S23 | Effect of DOTAP on MinD adsorption kinetics on DOPC and DOPG-DOPC SLBs | 24 |
| Figure S24 | Effect of DOTAP on MinD adsorption kinetics in DOPE and DOPG-DOPE SLBs | 25 |
| Figure S25 | Number of oscillations in the first 2h in SLBs with added DOTAP | 26 |
| Figure S26 | Effect of P70a-*minE* concentration on MinDE adsorption kinetics on a DOPG-DOPE SLB | 27 |
| Figure S27 | Effect of P70a-*minE* concentration on MinDE adsorption kinetics on a DOPG-DOPC SLB | 28 |
| Figure S28 | Effect of P70a-*minE* concentration on MinDE adsorption kinetics on an ECL SLB | 29 |
| Figure S29 | Tagging of Zorya proteins disrupts *E. coli* defense against phages | 30 |
| Figure S30 | Effect of DOPG on ZorA TXTL adsorption kinetics on a DOPE SLB | 31 |
| Figure S31 | Comparison of ZorA adsorption kinetics with a DOPC and a DOPE SLB | 32 |
| Figure S32 | ZorAB TXTL adsorption kinetics with an ECL SLB | 33 |
| Figure S33 | ZorE adsorption kinetics with post-incubation Tris-NaCl flush included in the graph | 34 |
| Table S1 | Working lipid concentrations during step III of SALB | 35 |

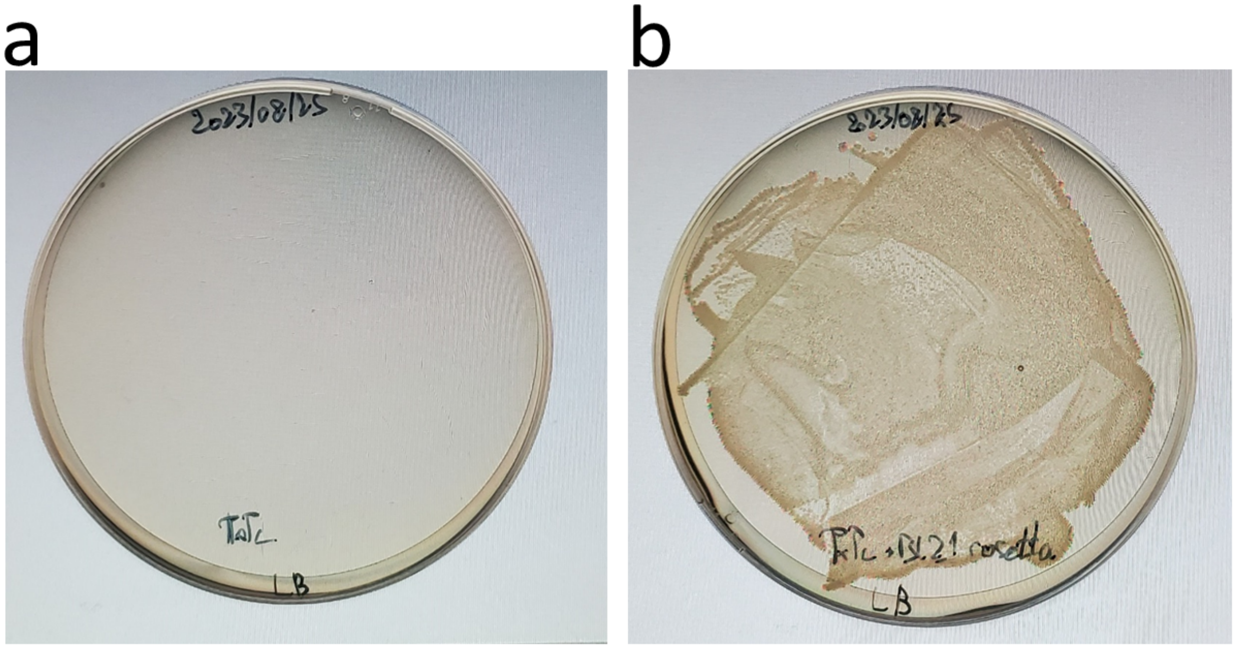


**Supplementary Figure 1. (a)** Plating 45 µL of a TXTL reaction onto an agar plate with no antibiotic shows no *E. coli* colonies. **(b)** Adding 5 µL of BL21 Rosetta *E. coli* to 40 µL of a TXTL reaction and then plating the mix onto an agar plate with no antibiotic shows many *E. coli* colonies.

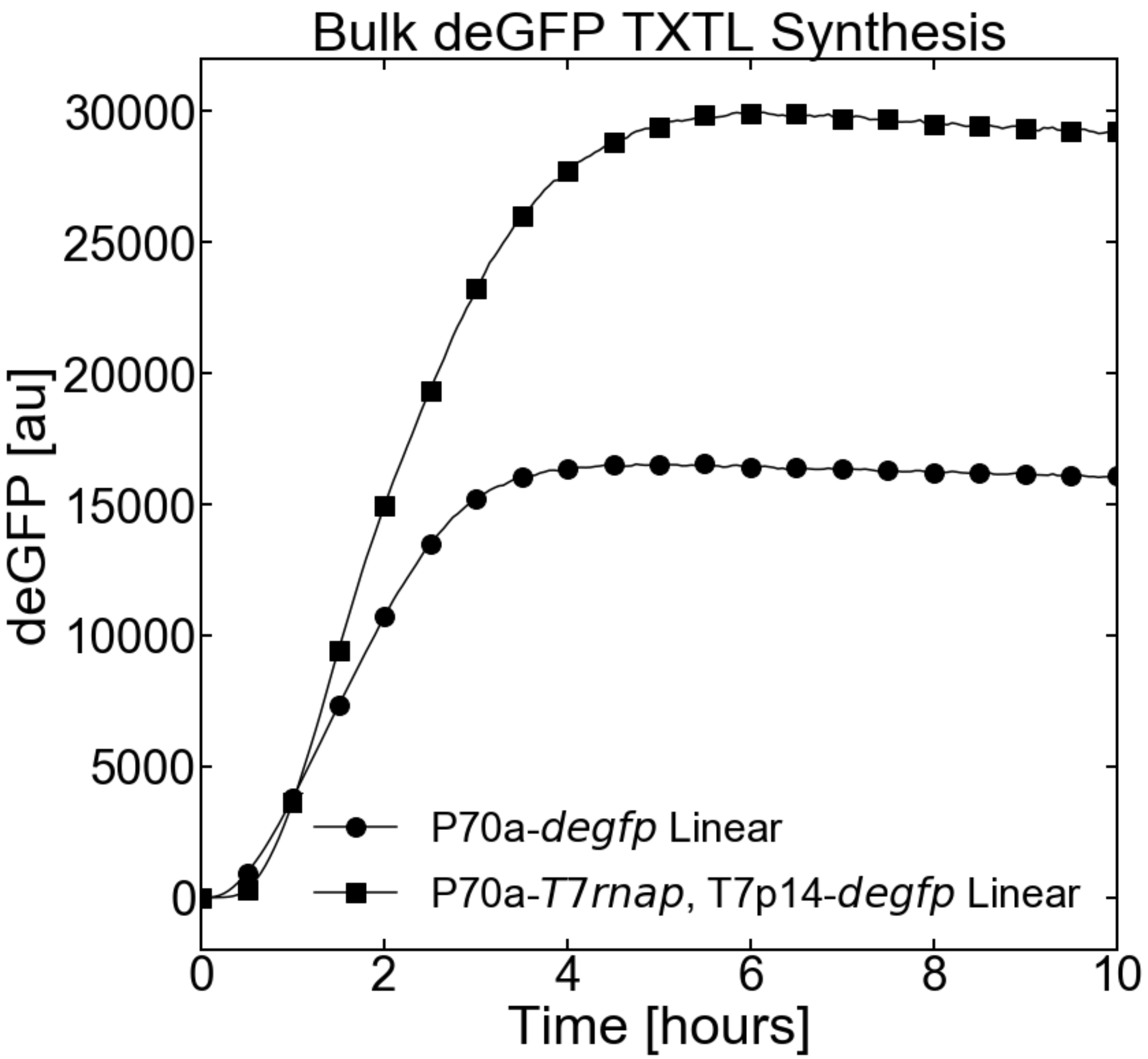


**Supplementary Figure 2.** Batch mode TXTL synthesis of deGFP with the P70a promoter (P70a-*degfp*, 5nM) and with the T7 cascade (P70a-*T7rnap*, 0.15nM, T7p14-*degfp*, 5nM).

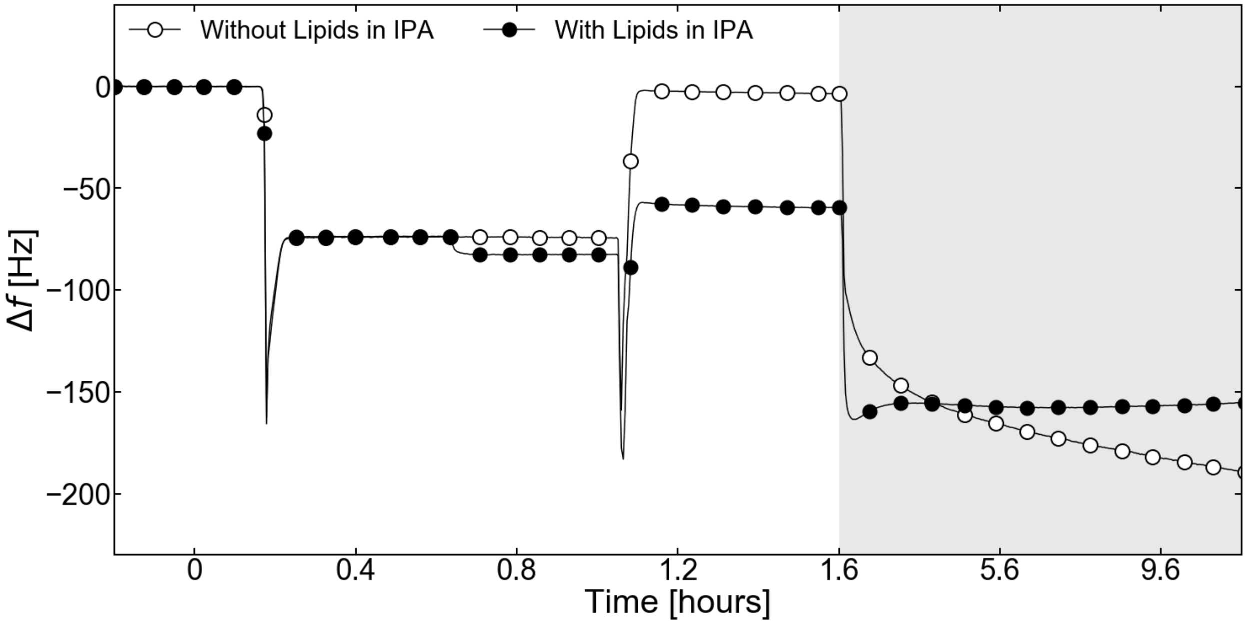


**Supplementary Figure 3.** SALB protocol completed with EggPC or without lipids added to the IPA-Lipid mix. Without phospholipids, step III shows a return to Δf = 0 Hz, which indicates no SLB has been formed.

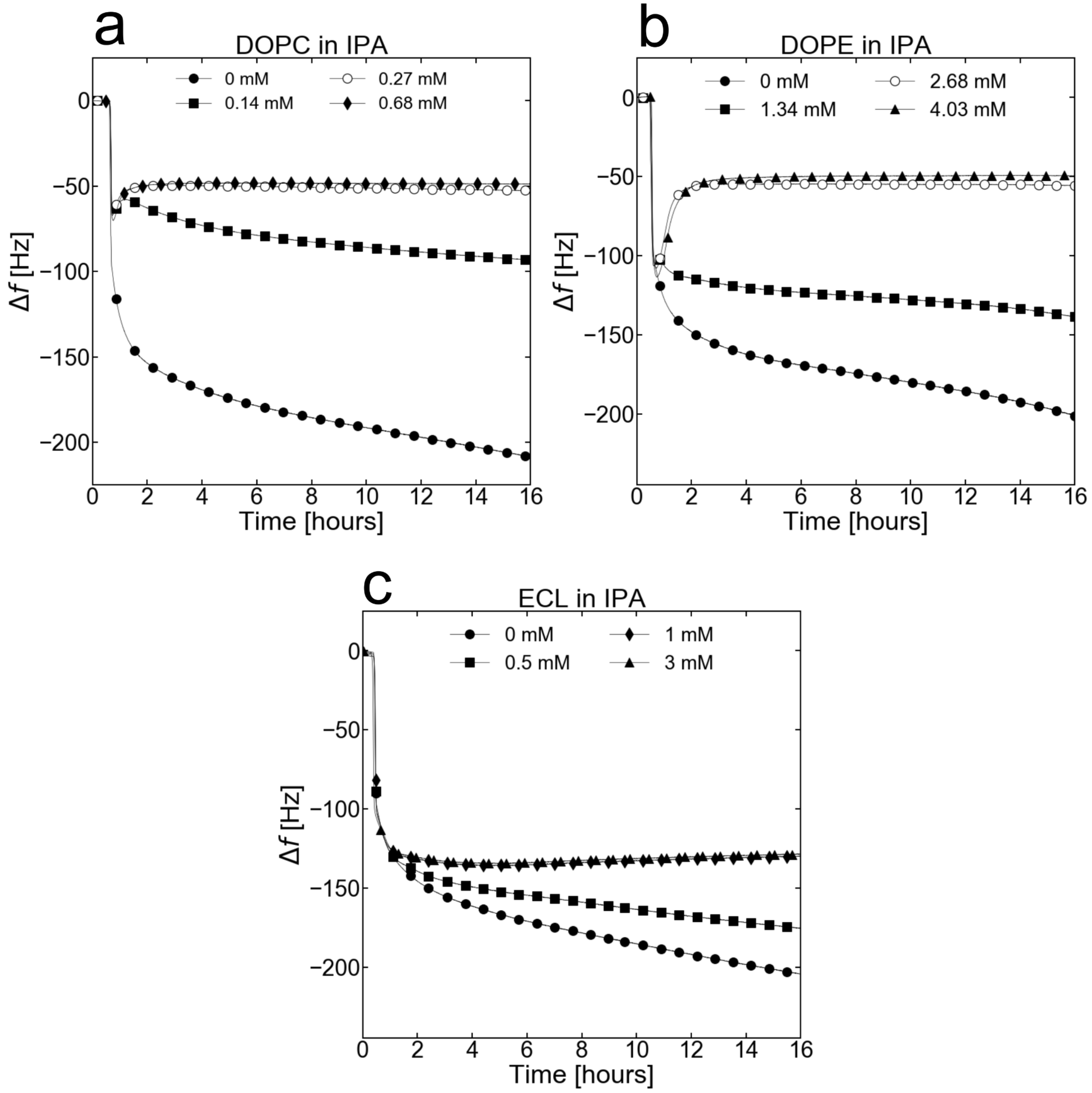


**Supplementary Figure 4.** Adsorption kinetics of a blank TXTL reaction (P70a-*T7rnap*, 0.15 nM) on SLBs depending on the DOPC **(a)**, DOPE **(b)**, and ECL **(c)** phospholipids concentration in IPA during Step II of SALB.

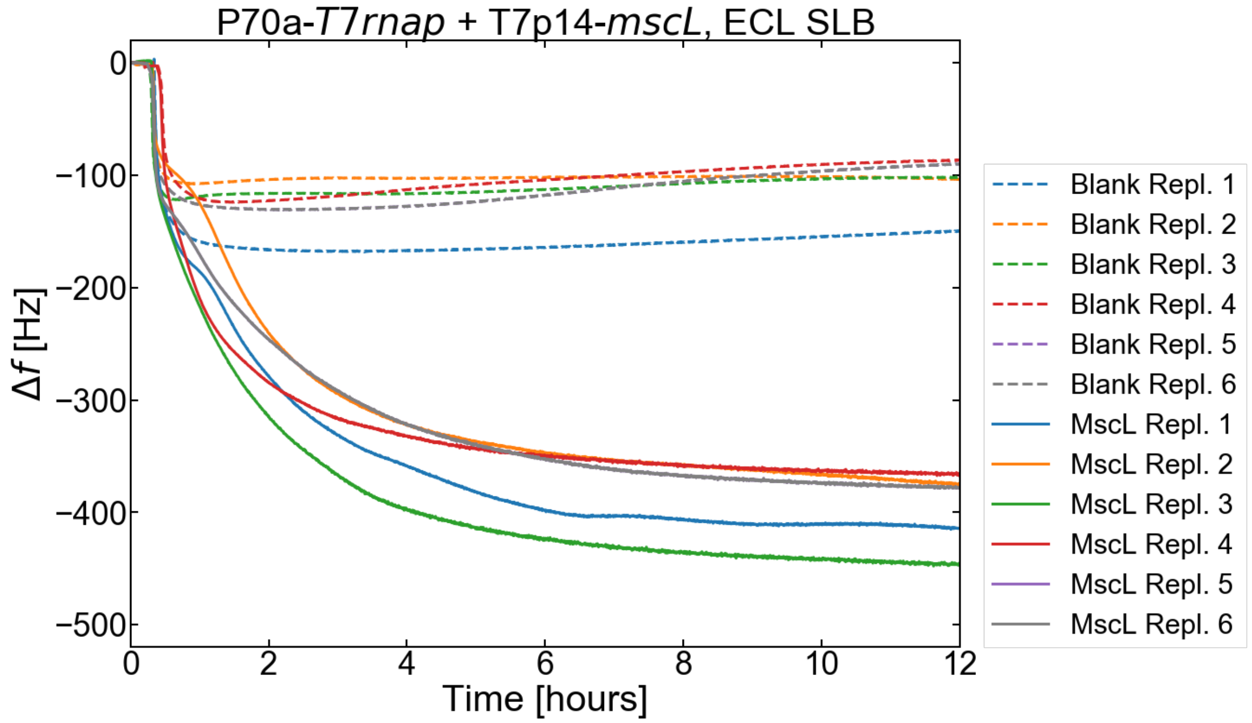


**Supplementary Figure 5.** Biological replicates of the MscL synthesizing TXTL reaction (P70a-*T7rnap*, 0.15 nM, T7p14-*mscL,* 5 nM) incubated on an ECL SLB.

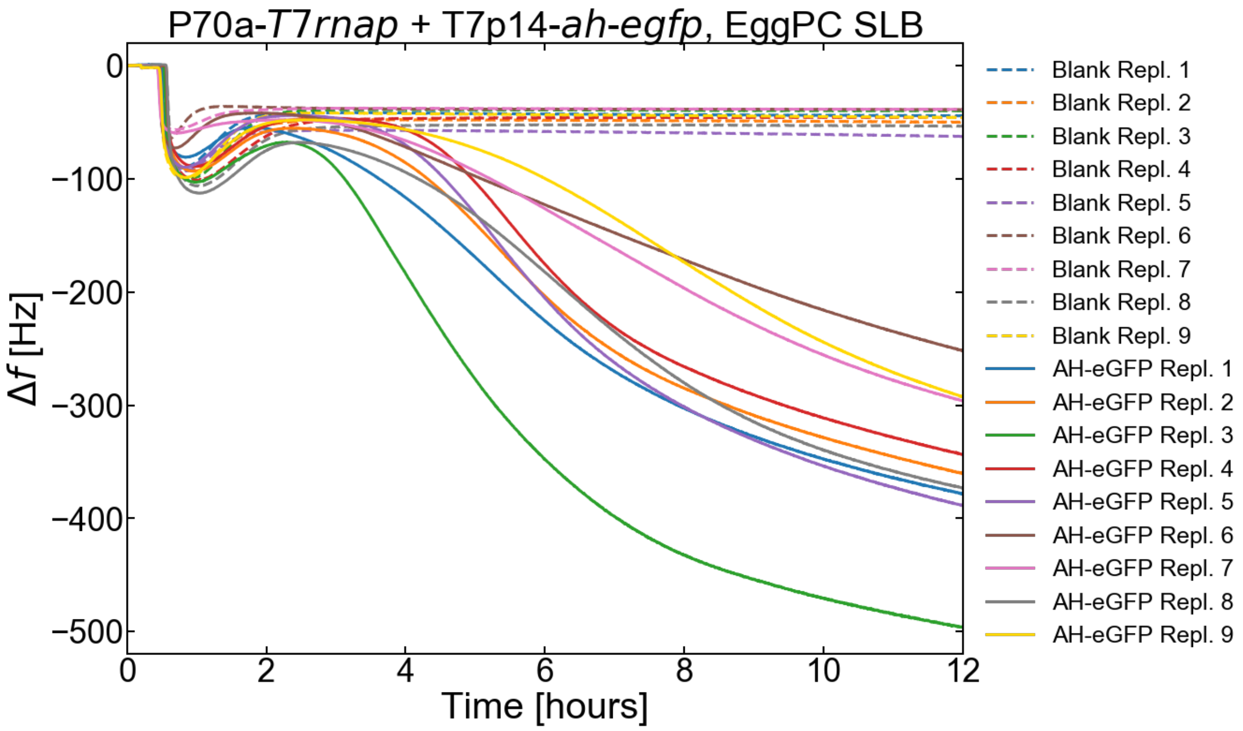


**Supplementary Figure 6.** Biological replicates of the AH-eGFP synthesizing TXTL reaction (P70a-*T7rnap*, 0.15 nM, T7p14-*ah-egfp,* 5 nM) incubated on an EggPC SLB.

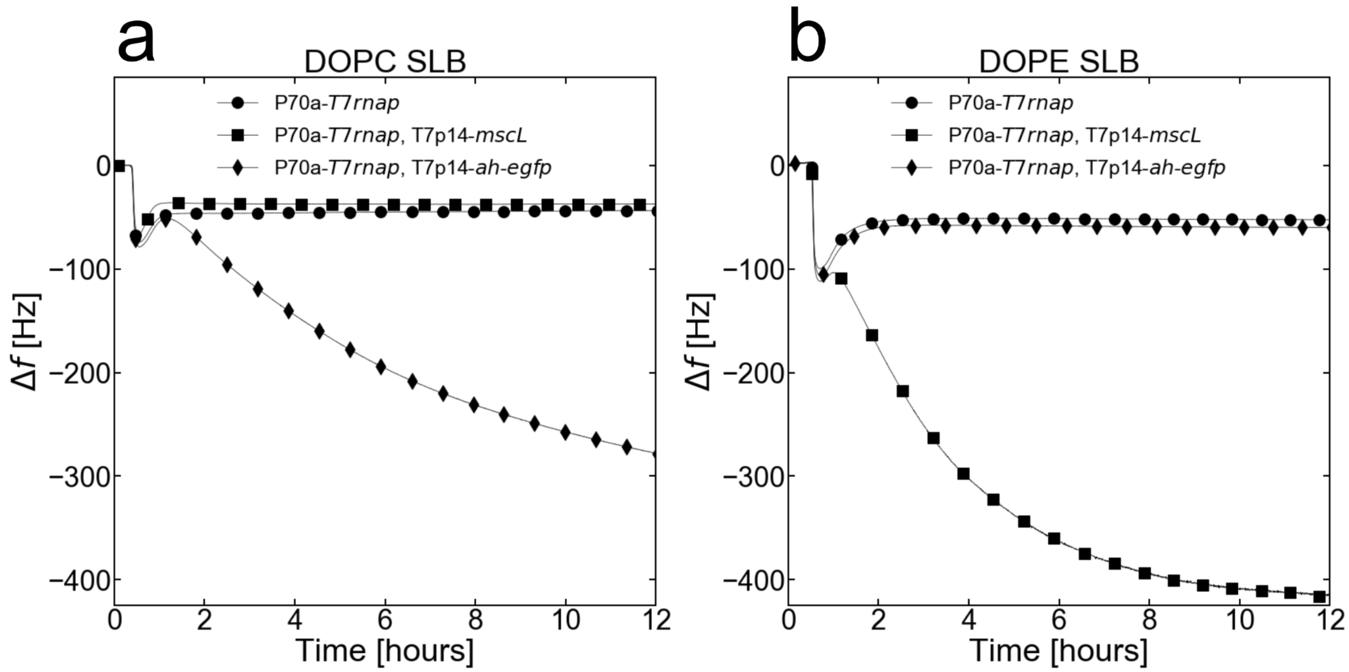


**Supplementary Figure 7. (a) and (b)** Adsorption kinetics of either blank (P70a-*T7rnap*, 0.15 nM), AH-eGFP (P70a-*T7rnap*, 0.15 nM, T7p14-*ah-egfp,* 5 nM), or MscL (P70a-*T7rnap*, 0.15 nM, T7p14-*mscL,* 5 nM) TXTL reactions into a DOPC SLB and a DOPE SLB respectively.

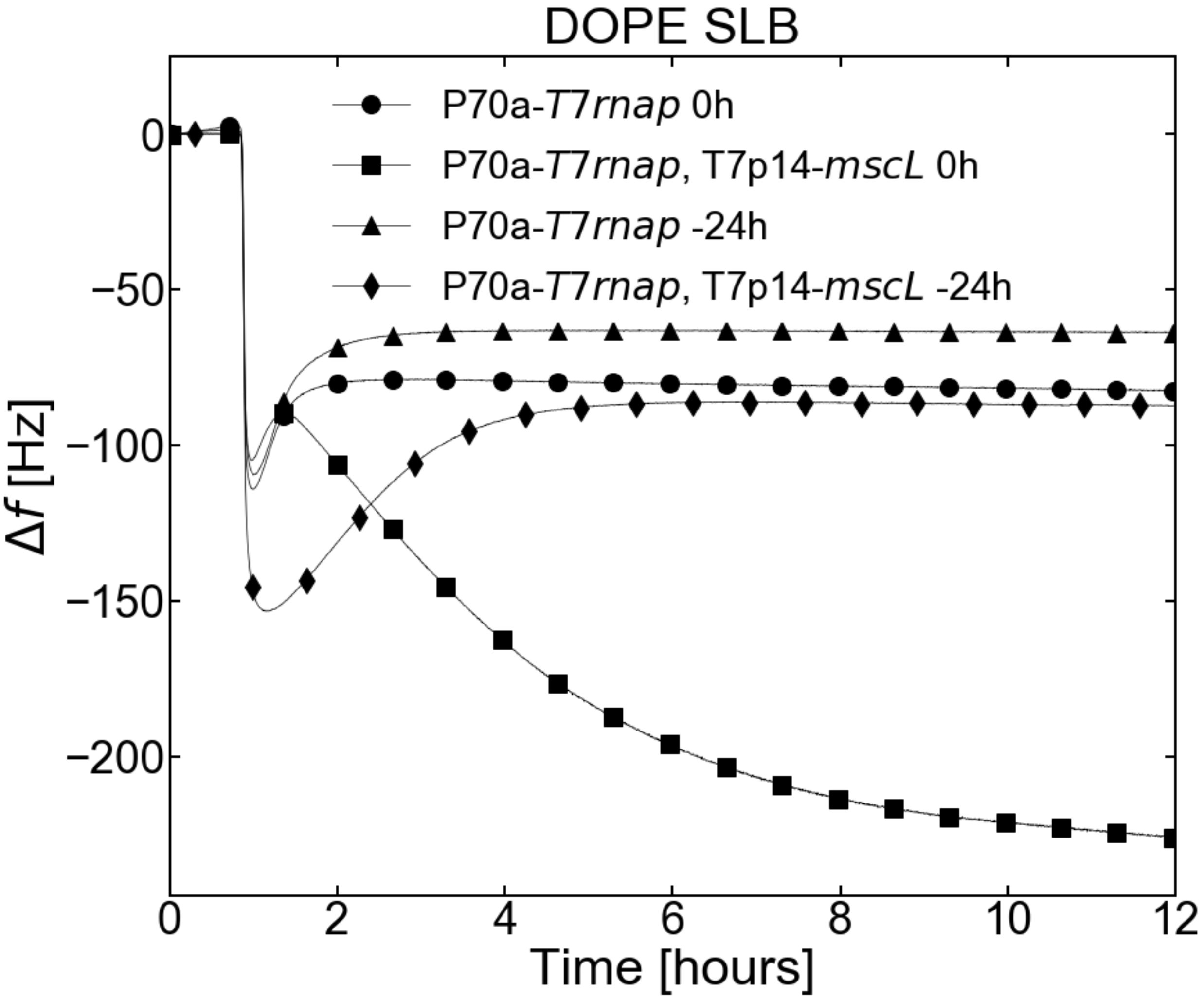


**Supplementary Figure 8.** Adsorption kinetics into a pure DOPE SLB of a blank TXTL (P70a-*T7rnap*, 0.15 nM) and an MscL TXTL (P70a-*T7rnap*, 0.15 nM, T7p14-*mscL,* 5 nM) reactions that have either been freshly mixed (0 h) or that have been pre-incubated for 24 h (-24 h) before being flushed into the QCMD module.

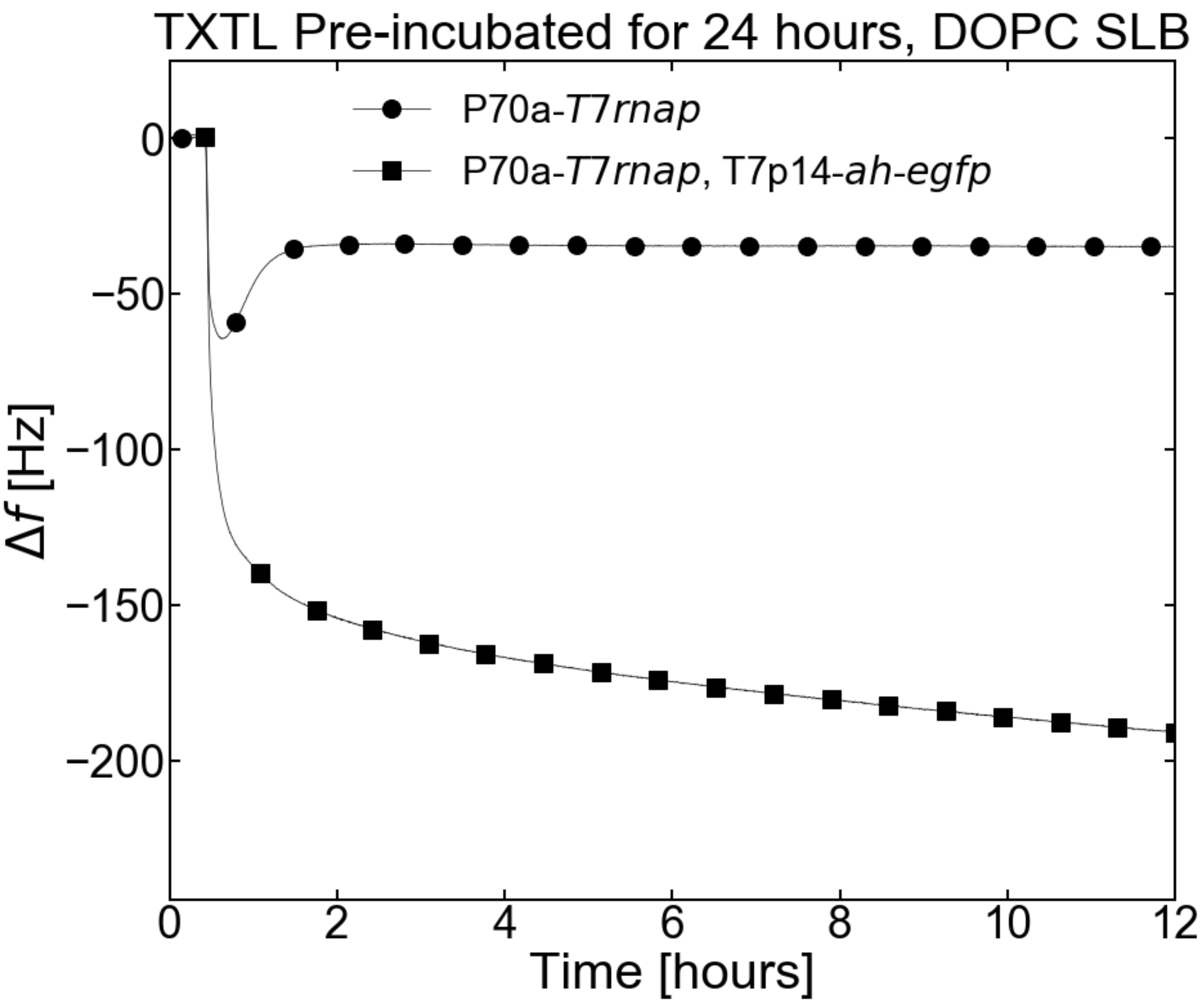


**Supplementary Figure 9.** Adsorption kinetics onto a pure DOPC SLB of a blank TXTL (P70a-*T7rnap*, 0.15 nM) and an AH-eGFP TXTL (P70a-*T7rnap*, 0.15 nM, T7p14-*ah-egfp,* 5 nM), reactions that have been pre-incubated for 24 h before being flushed into the QCMD module.

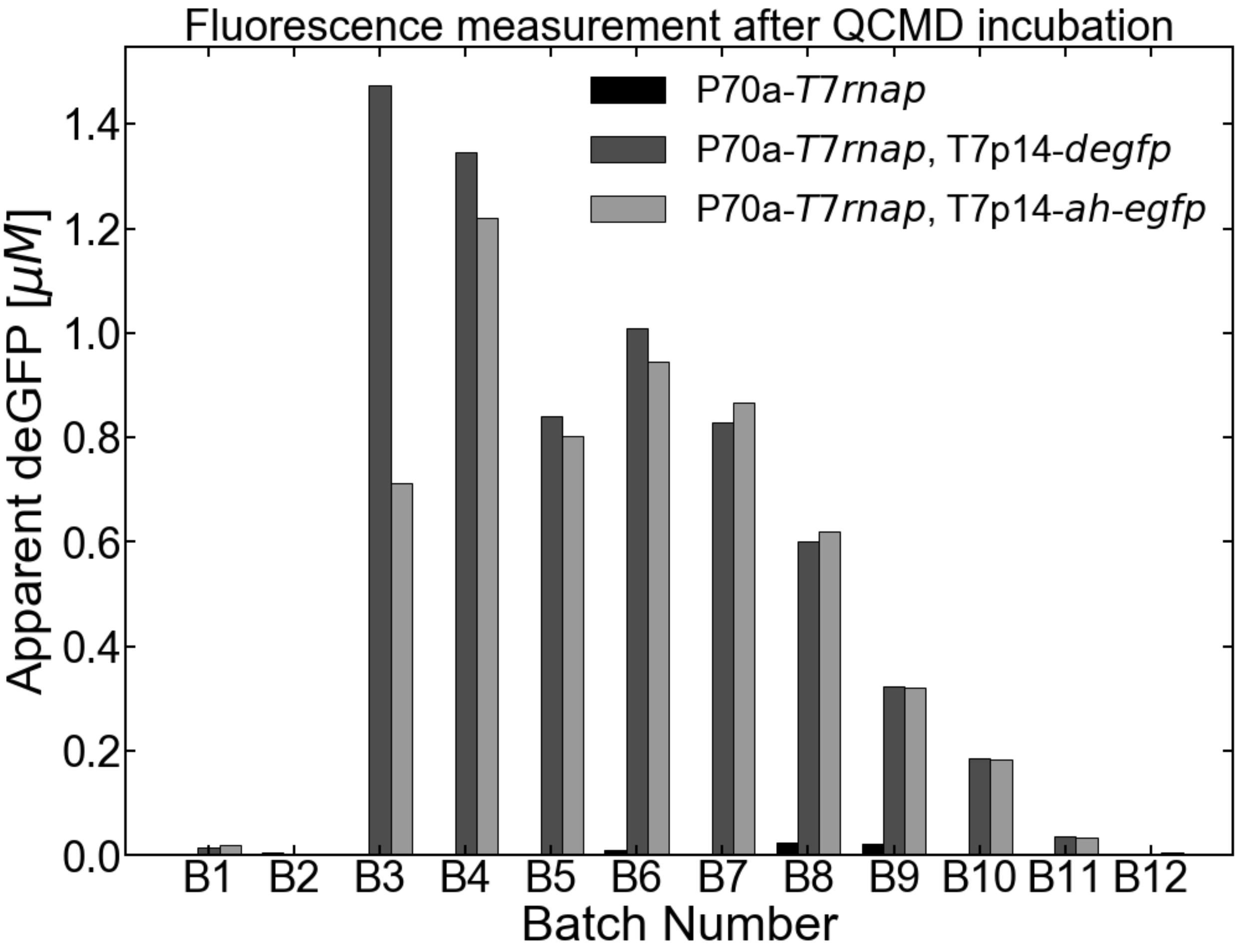


.
**Supplementary Figure 10.** End-point measurement of either blank (P70a-*T7rnap*, 0.15 nM), AH-eGFP (P70a-*T7rnap*, 0.15 nM, T7p14-*ah-egfp,* 5 nM), or MscL (P70a-*T7rnap*, 0.15 nM, T7p14-*mscL,* 5 nM) TXTL reactions in the QCMD modules overnight. The TXTL is flushed out of the module in batches of 100 µL and the x-axis corresponds to the number of the batch.

| **Sensor dimensions** |
|---|
| - Diameter 1.2 cm<br>- Surface area = 1.131 $10^{-4}$ $m^2$ |
| **MscL dimensions** |
| - MscL assembles into a pentamer<br>- Molar mass of MscL monomer = 14957.2 g/mol<br>- Mass of one monomer = 2.48 $10^{-20}$ g<br>- Mass of one pentamer = 1.24 $10^{-19}$ g<br>- Surface area of one pentamer: 140 $nm^2$ [1] |
| **Mass of closely packed MscL pentamers on the SLB** |
| Hypothesis: close packing of circular MscL pentamers on SLB:<br>- The size ratio of closely packed circles is about 0.6<br>- The number of MscL pentamers on the SLB is: 0.6 * 1.131 $10^{-4}$ / (1.4 $10^{-16}$) = 4.85 $10^{11}$<br>- The maximum total mass of MscL on the SLB is 60.1 ng<br>- For a 40-µl reaction (volume of each QCMD chamber), this corresponds to a concentration of 1.5 µg/ml or 0.1 µM of MscL proteins. Therefore, based on our deGFP and AH-eGFP quantifications (1-2 µM produced in QCMD chambers), about 20 times more proteins are produced in the QCMD chamber with respect to the membrane capacity<br>- We assume a resolution of 5 Hz from the QCMD frequency signal<br>- For MscL the drop is of about 300 Hz<br>- We can detect 60 times less MscL, which corresponds to about 1 ng, based on maximum membrane coverage with MscL<br>- It is unlikely that MscL reaches this level of packing on our SLBs, so this mass sensitivity is likely to be underestimated |

**Supplementary Figure 11.** Estimation of the mass sensitivity of the QCMD.

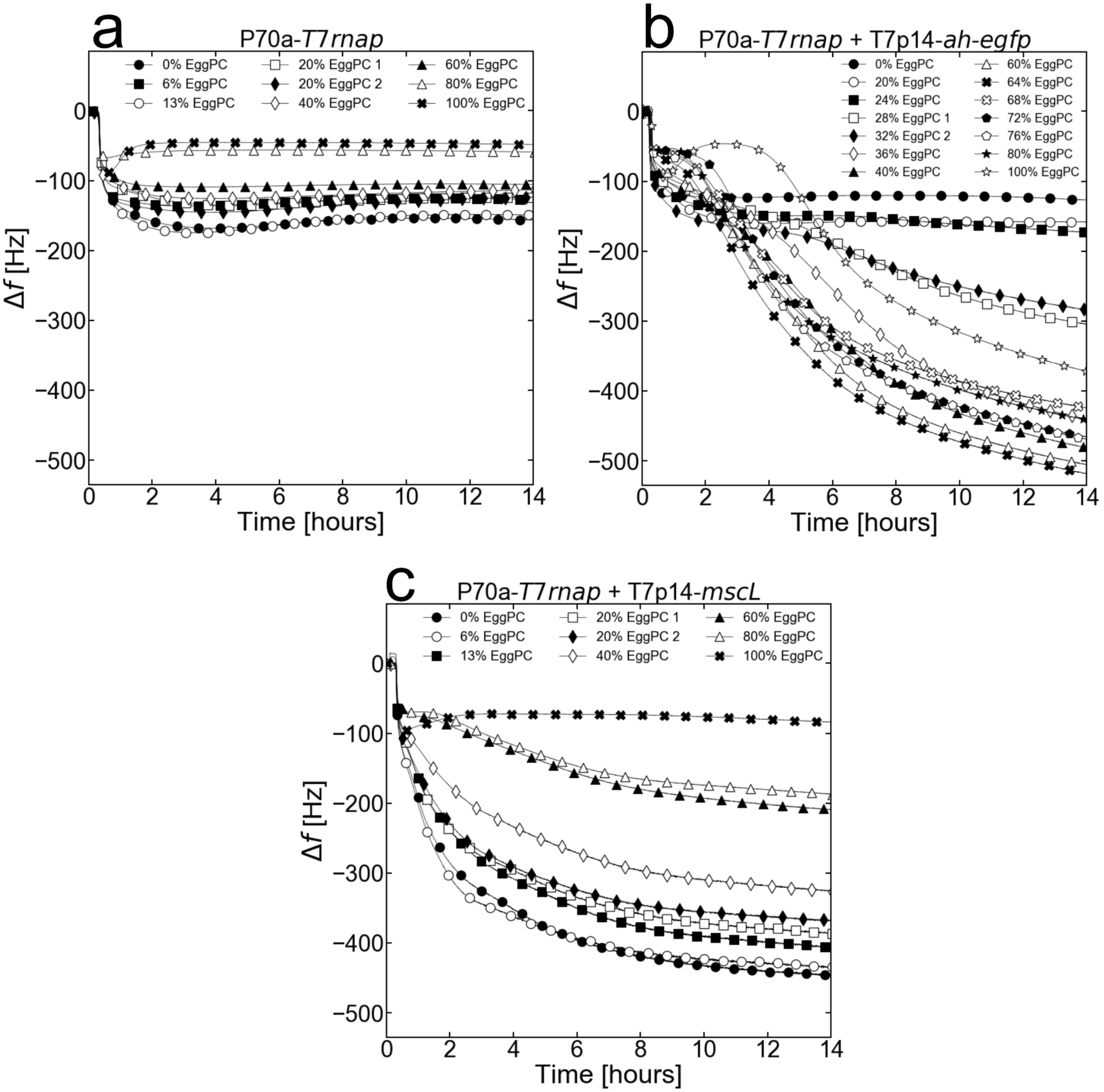


**Supplementary Figure 12.** Adsorption kinetics of **(a)** a blank (P70a-*T7rnap*, 0.15 nM), **(b)** an AH-eGFP (P70a-*T7rnap*, 0.15 nM, T7p14-*ah-egfp*, 5 nM), and **(c)** a MscL (P70a-*T7rnap*, 0.15 nM, T7p14-*mscL*, 5 nM) TXTL reaction respectively into ECL – EggPC hybrid SLBs.

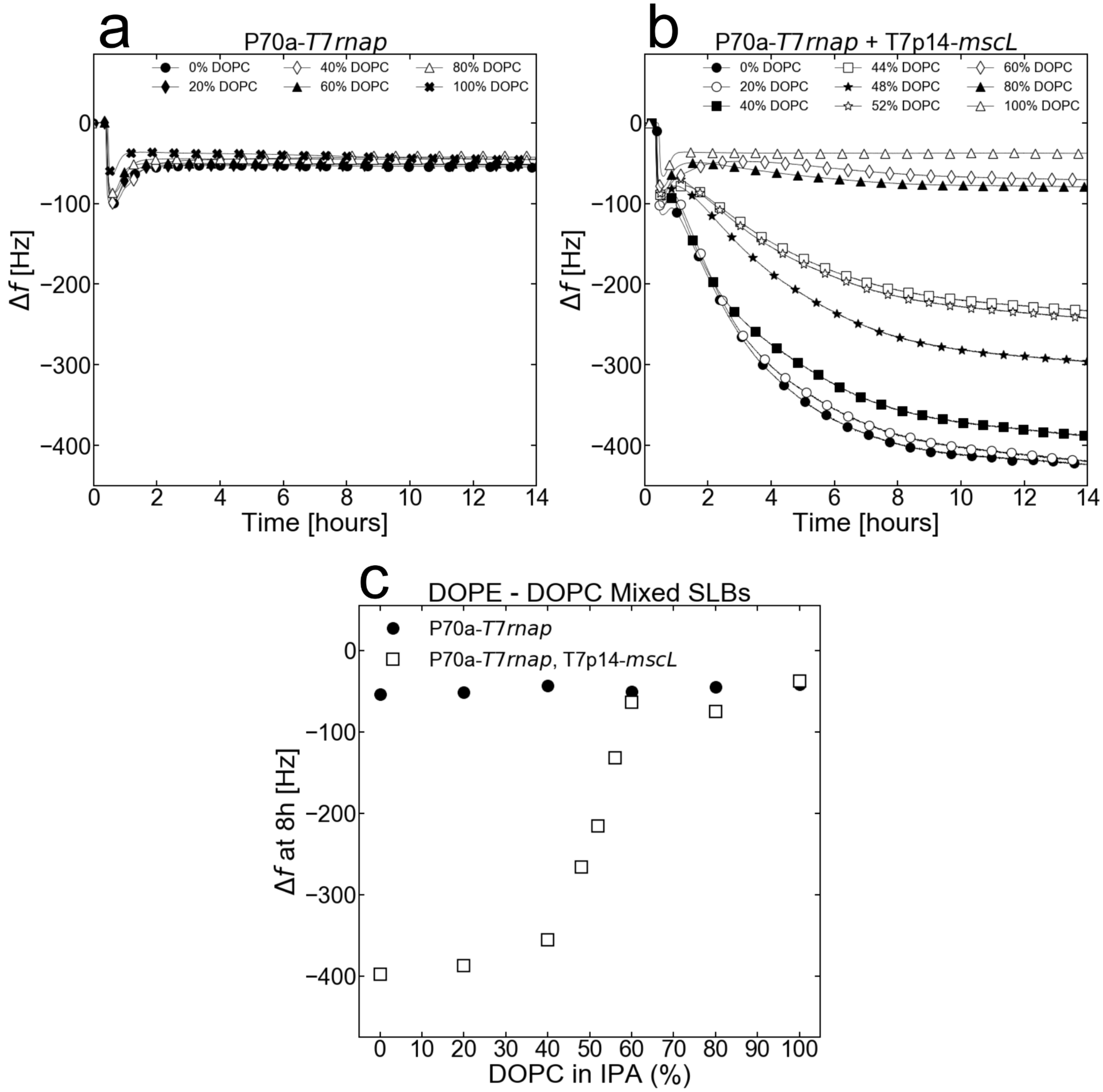


**Supplementary Figure 13. (a) and (b)** Adsorption kinetics of a blank (P70a-*T7rnap*, 0.15 nM) and a MscL (P70a-*T7rnap*, 0.15 nM, T7p14-*mscL*, 5 nM) TXTL reactions respectively into DOPE – DOPC Mixed SLBs. **(c)** Frequency changes after 8 hours of incubating either blank (P70a-*T7rnap*, 0.15 nM) or MscL (P70a-*T7rnap*, 0.15 nM, T7p14-*mscL*, 5 nM) TXTL reactions into DOPE – DOPC Mixed SLBs.

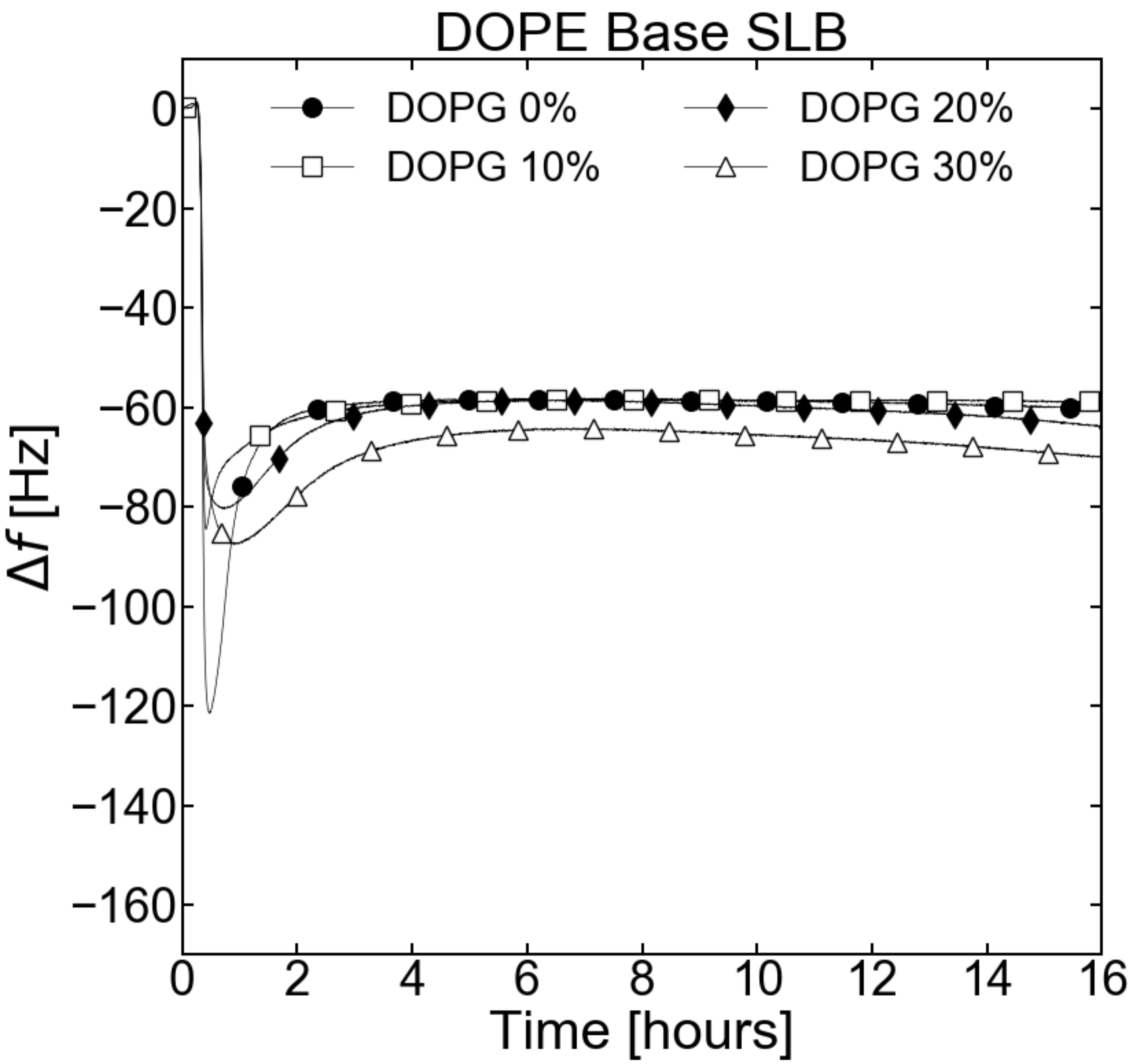


**Supplementary Figure 14.** Adsorption kinetics of a blank TXTL reaction (P70a-*T7rnap*, 0.15 nM) onto 0, 10, 20, and 30% DOPG/DOPE SLBs

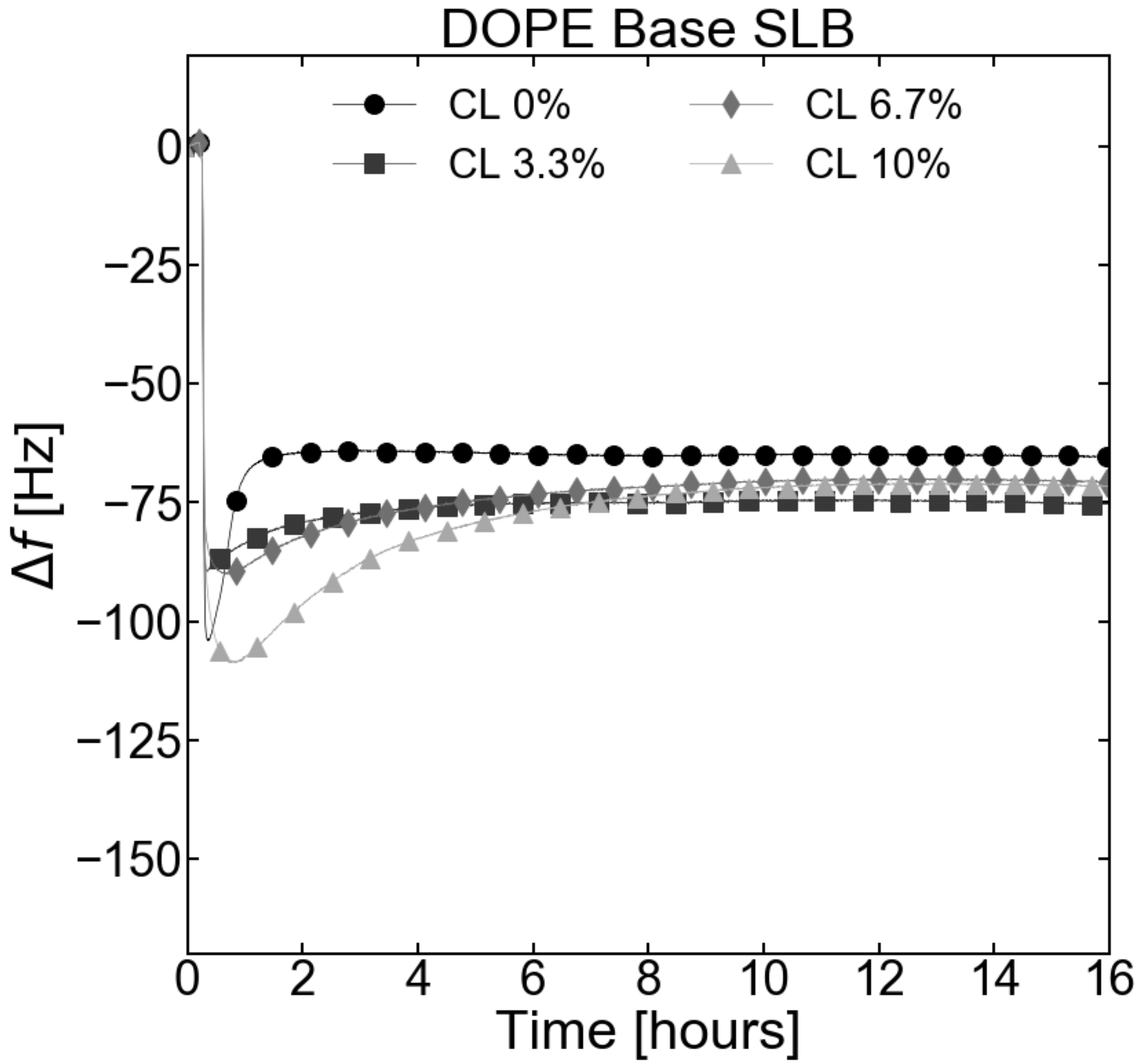


**Supplementary Figure 15.** Adsorption kinetics of a blank TXTL reaction (P70a-*T7rnap*, 0.15 nM) onto 0, 3.3, 6.7, and 10% CL/DOPE SLBs.

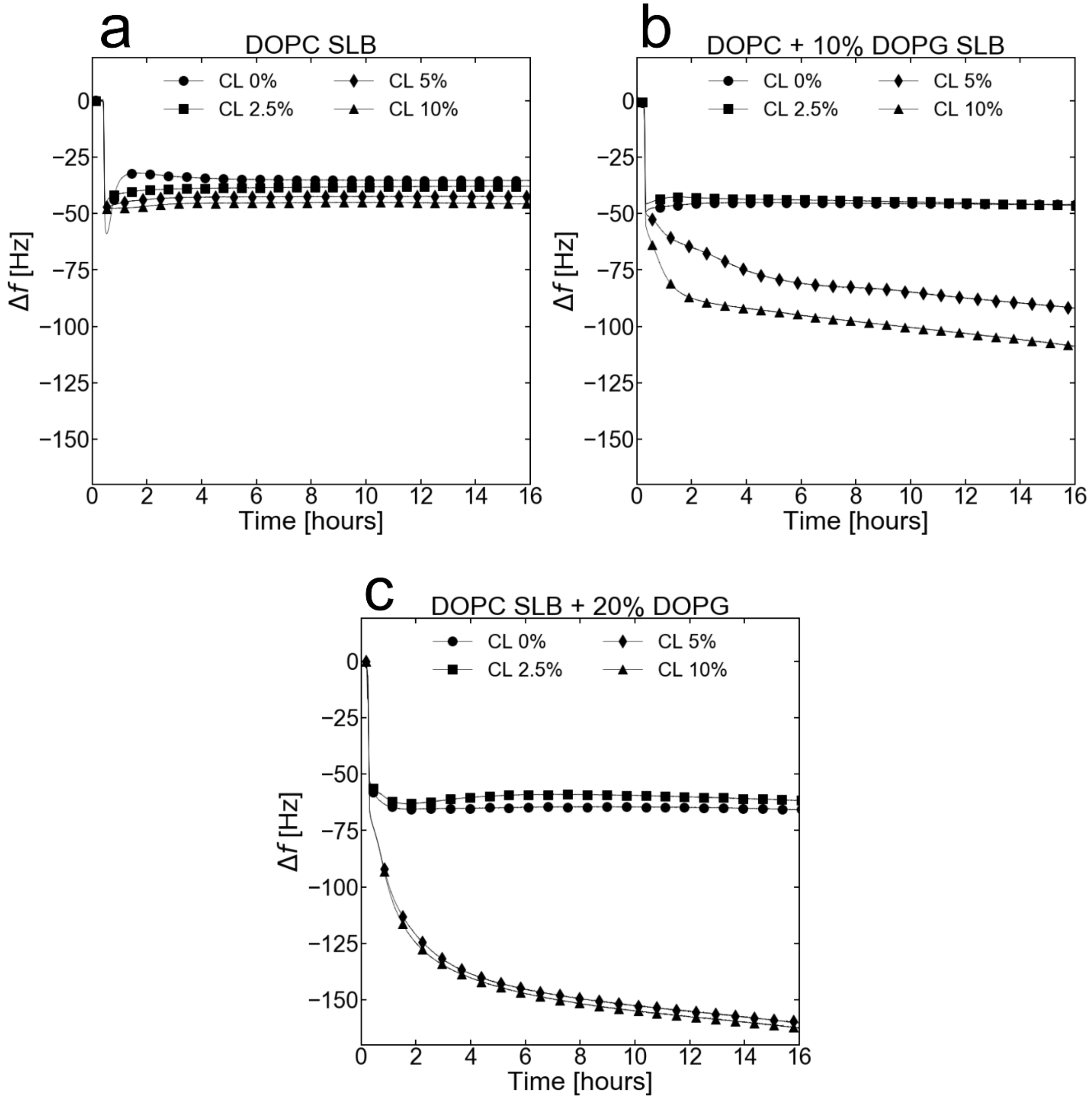


**Supplementary Figure 16. (a), (b), and (c)** Adsorption kinetics of a blank TXTL reaction (P70a-*T7rnap*, 0.15 nM) onto a DOPC Base SLB as for different relative CL concentrations for a 0, 10, and 20% DOPG/DOPC SLBs respectively.

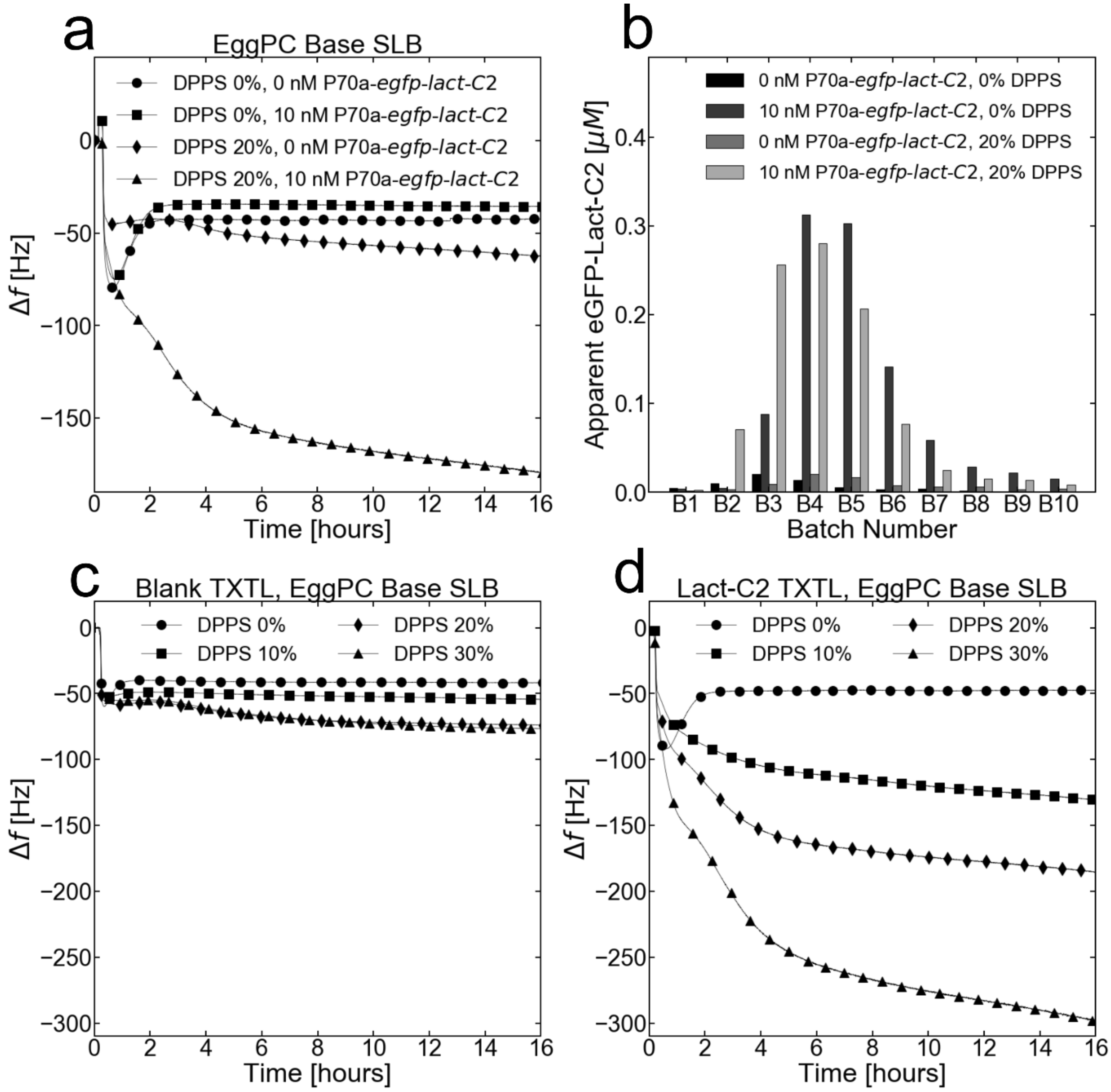


**Supplementary Figure 17. (a)** Adsorption kinetics of either a blank (no DNA) or eGFP-Lact-C2 (P70a-*egfp-lact-C2*) TXTL reactions incubated onto either a pure EggPC SLB or an EggPC SLB mixed with DPPS. **(b)** End-point measurement of TXTL expression of panel **(a)**. The TXTL is flushed out of the module in batches of 100 μL and the x-axis corresponds to the number of the batch. **(c)** Adsorption kinetics of a blank TXTL (no DNA added) reaction incubated onto a 0, 10, 20, and 30% DPPS/EggPC mol. Ratio SLBs. **(d)** Adsorption kinetics of an eGFP-Lact-C2 TXTL (P70a-*egfp-lact-C2*, 10 nM) reaction incubated onto a 0, 10, 20, and 30% DPPS/EggPC mol. Ratio SLBs.

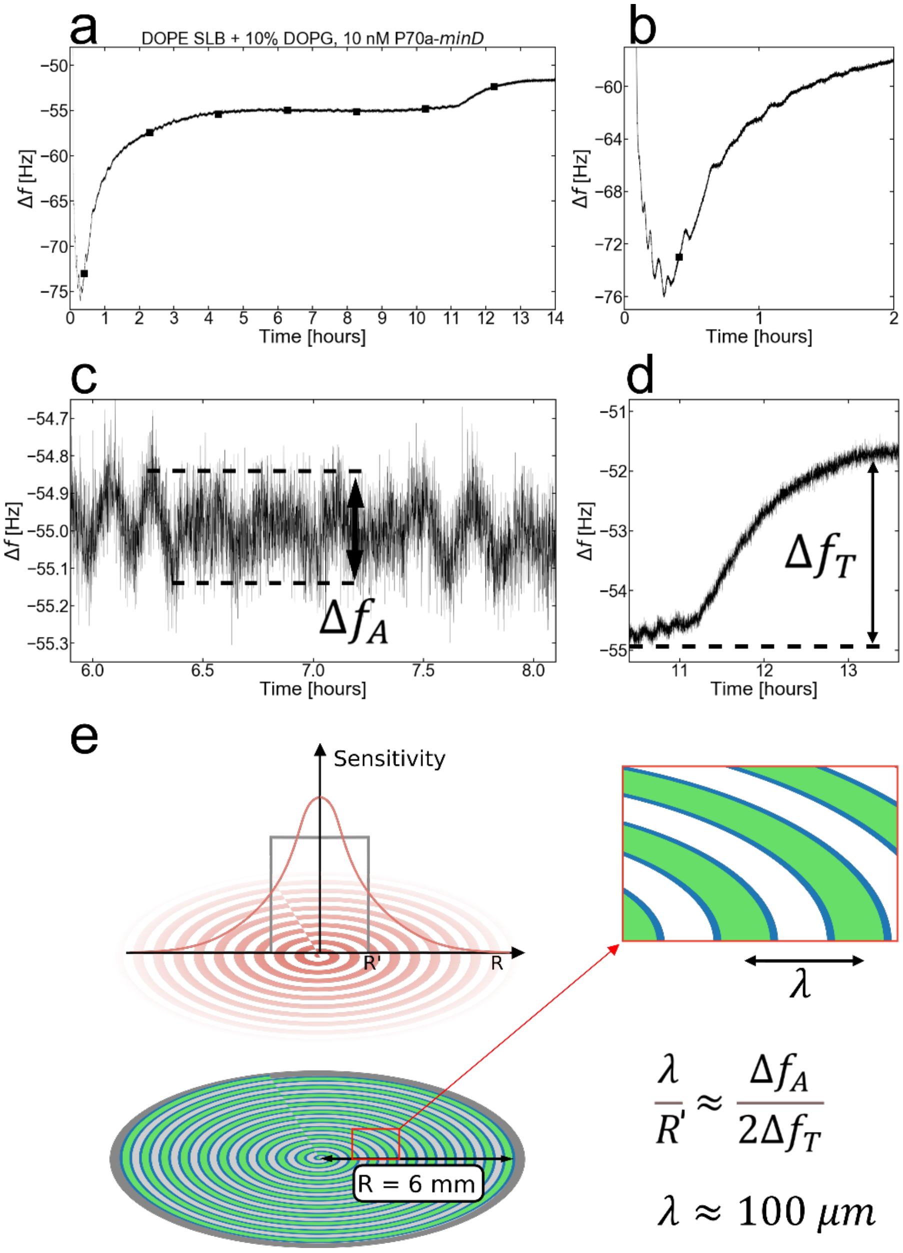


**Supplementary Figure 18.** **(a)** Adsorption kinetics of a MinD TXTL (P70a-*minD*, 10 nM) reaction incubated onto DOPE + 10% DOPG SLB. **(b)** Large amplitude oscillations during the first 2 hours of incubation. **(c)** Small-amplitude oscillations during the middle of the incubation. The amplitude of a single oscillation is labeled as $\Delta f_A$. **(d)** The increase in frequency at the end of the reaction due to ATP depletion is labeled as $\Delta f_T$. **(e)** Assuming that the Min patterns are radially symmetric fronts moving from the center of the sensor-SLB system to the edge, then $\Delta f_A$ is proportional to the mass of the outermost ring escaping the boundaries of the most sensitive area of the sensor (grey box, $r < R' \approx R/3)^2$, while $\Delta f_T$ is proportional to the mass of all rings combined within the most sensitive area of the sensor. Since the escaping ring is twice the mass of the average ring within the box, then we can approximate the wavelength as in the equation in the panel. This approximation yields a wavelength of approximately 100 μm from the parameters obtained in **(c)** and **(d)**.

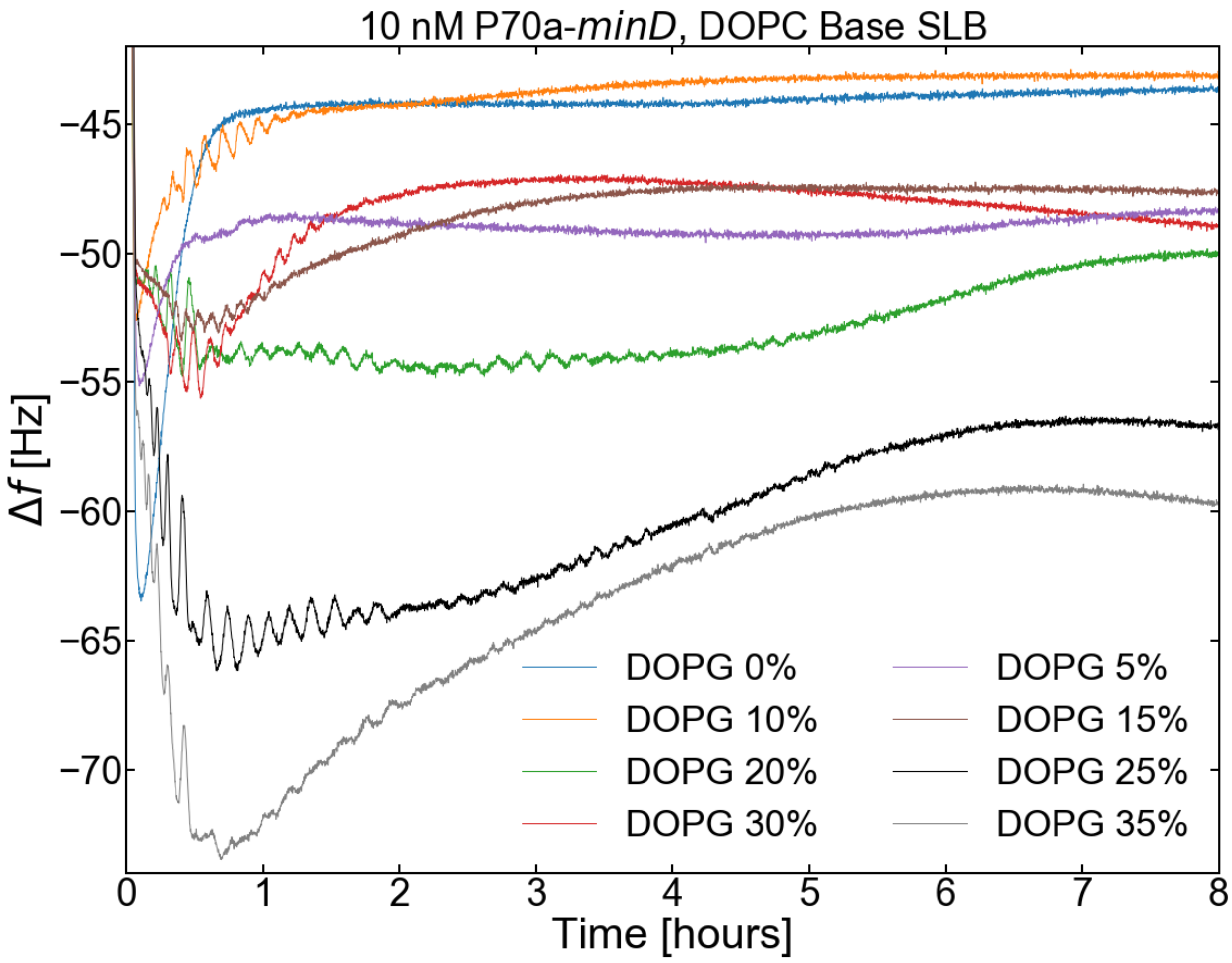


**Supplementary Figure 19.** Adsorption kinetics of a MinD TXTL (P70a-*minD*, 10 nM) reaction for different DOPG/DOPC SLBs.

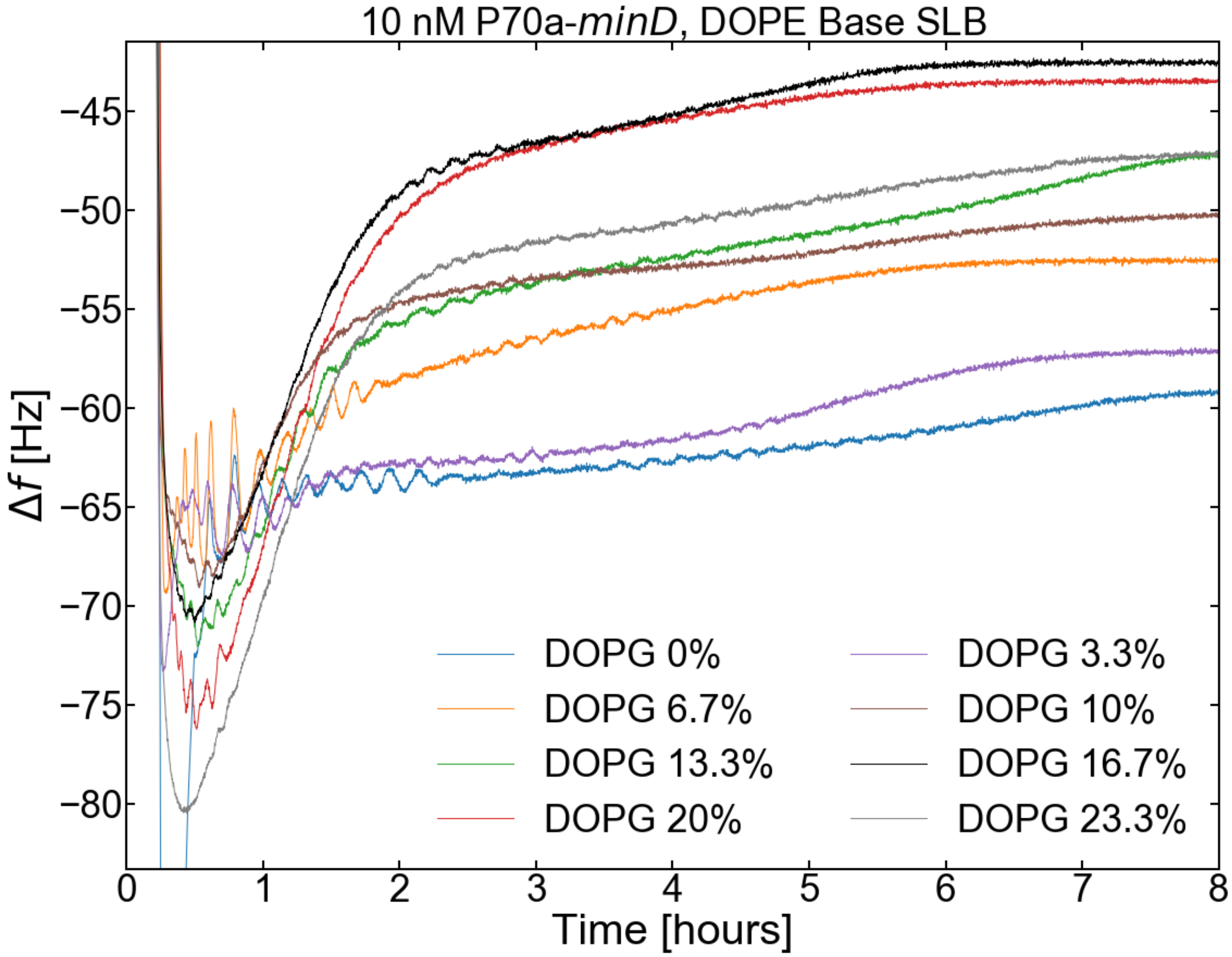


**Supplementary Figure 20. (a)** Adsorption kinetics of a MinD TXTL (P70a-*minD*, 10 nM) reaction for different DOPG/DOPE SLBs.

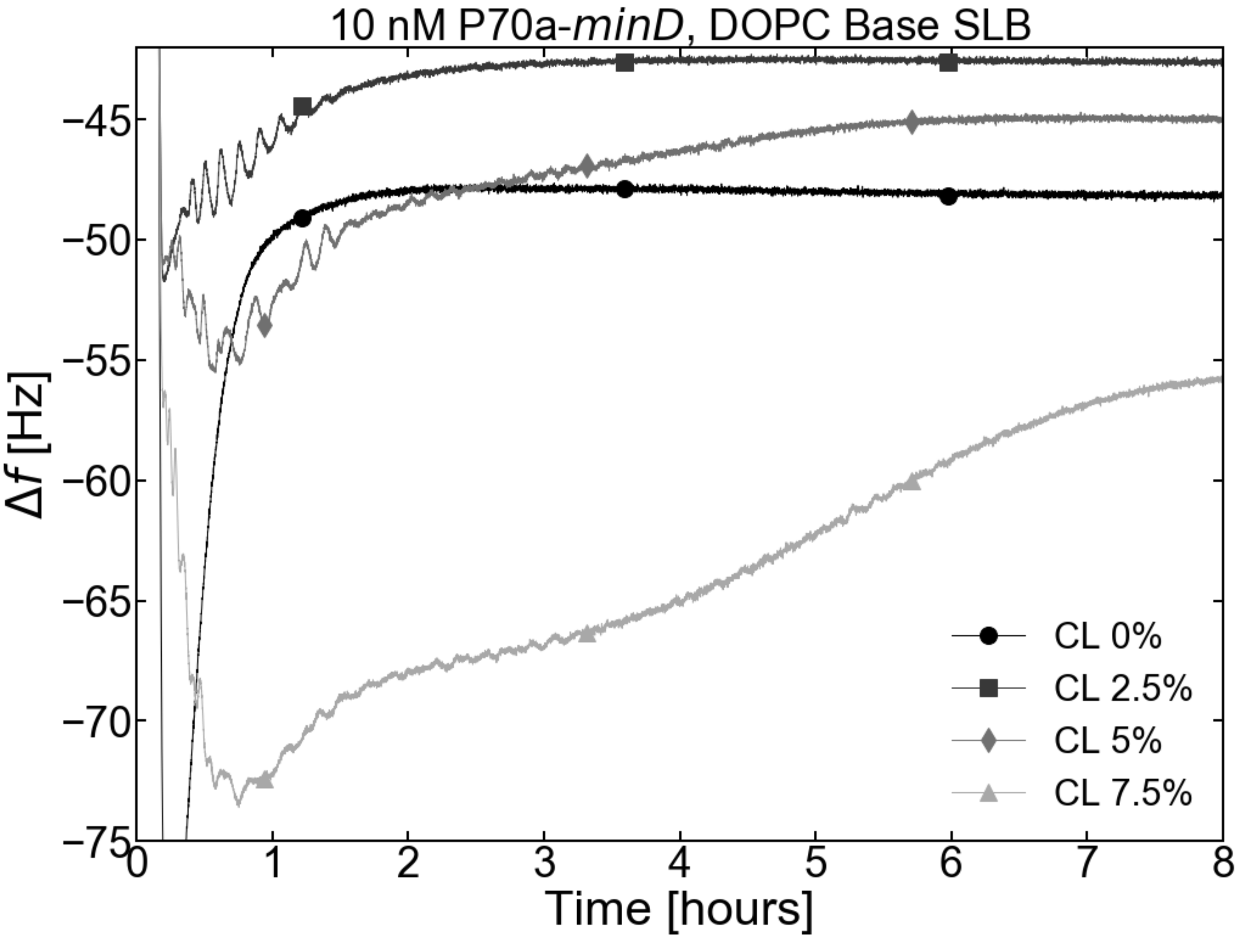


**Supplementary Figure 21.** Adsorption kinetics of a MinD TXTL (P70a-*minD*, 10 nM) reaction for different CL/DOPC SLBs.

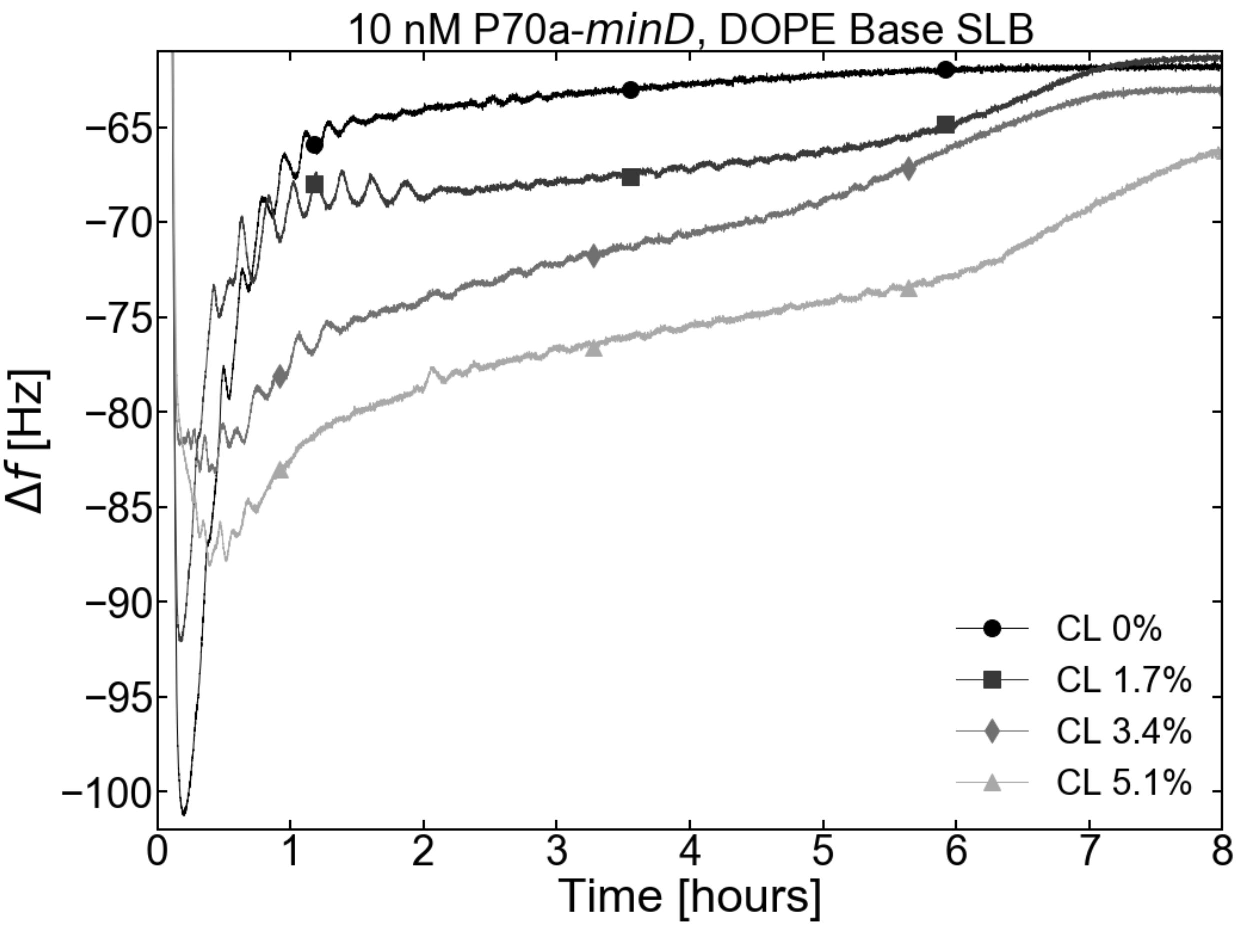


**Supplementary Figure 22.** Adsorption kinetics of a MinD TXTL reaction (P70a-*minD*, 10 nM) for different CL/DOPE SLBs.

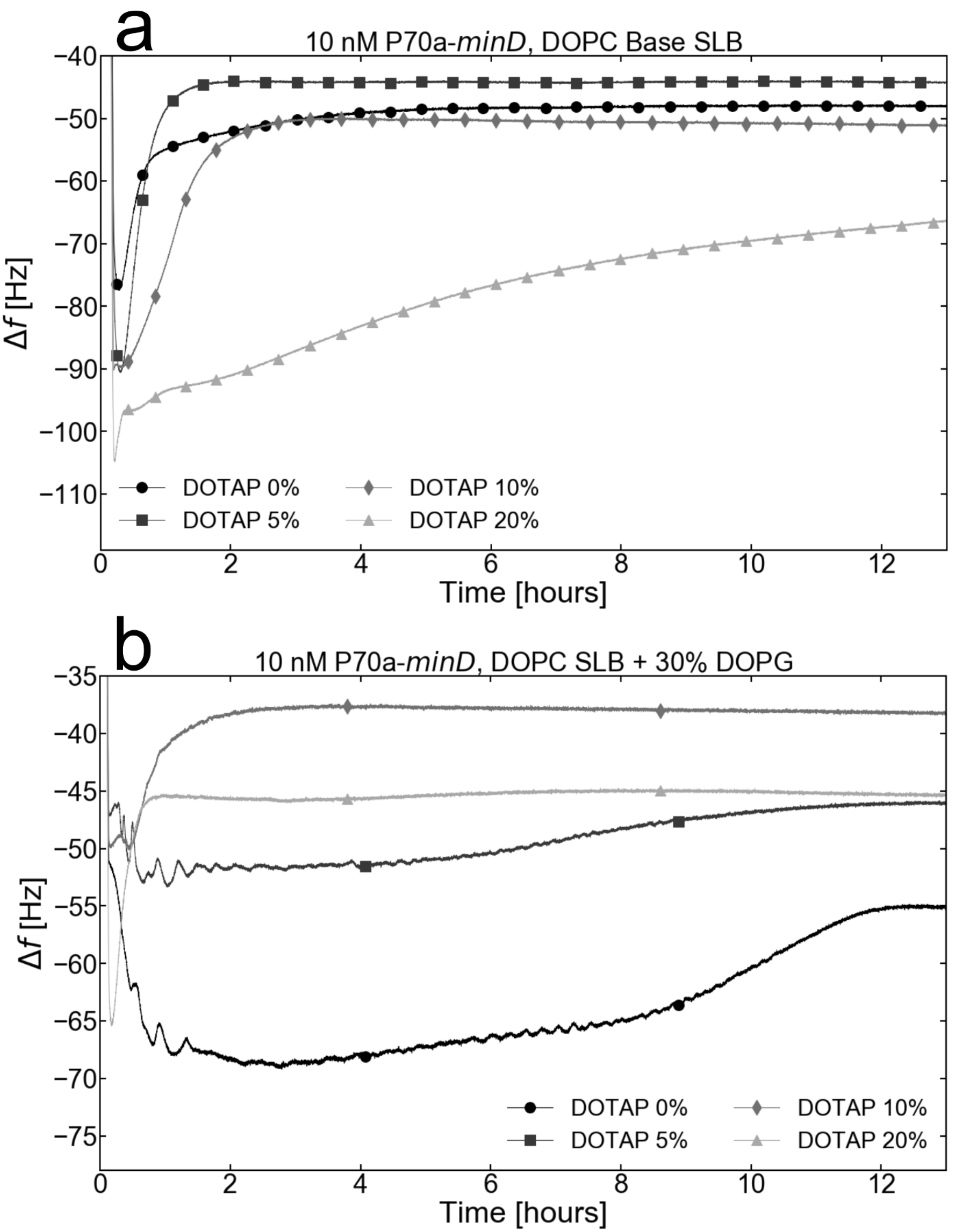


**Supplementary Figure 23. (a) and (b)** Adsorption kinetics of a MinD TXTL (P70a-*minD*, 10 nM) reaction for a range of different DOTAP/DOPC SLBs and DOTAP/DOPG/DOPC SLBs respectively.

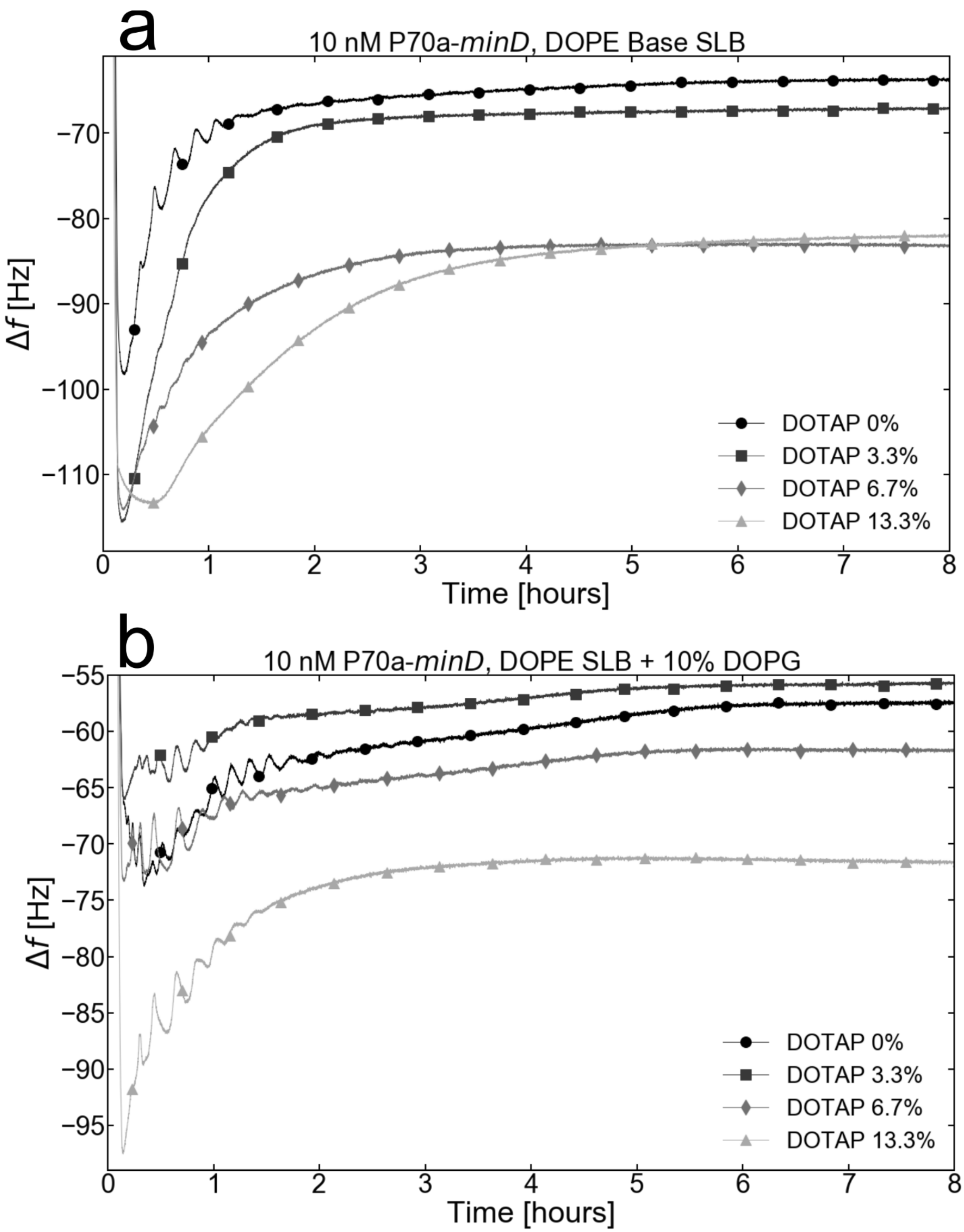


**Supplementary Figure 24. (a) and (b)** Adsorption kinetics of a MinD TXTL (P70a-*minD*, 10 nM) reaction for a range of different DOTAP/DOPE SLB and DOTAP/DOPG/DOPE SLB respectively.

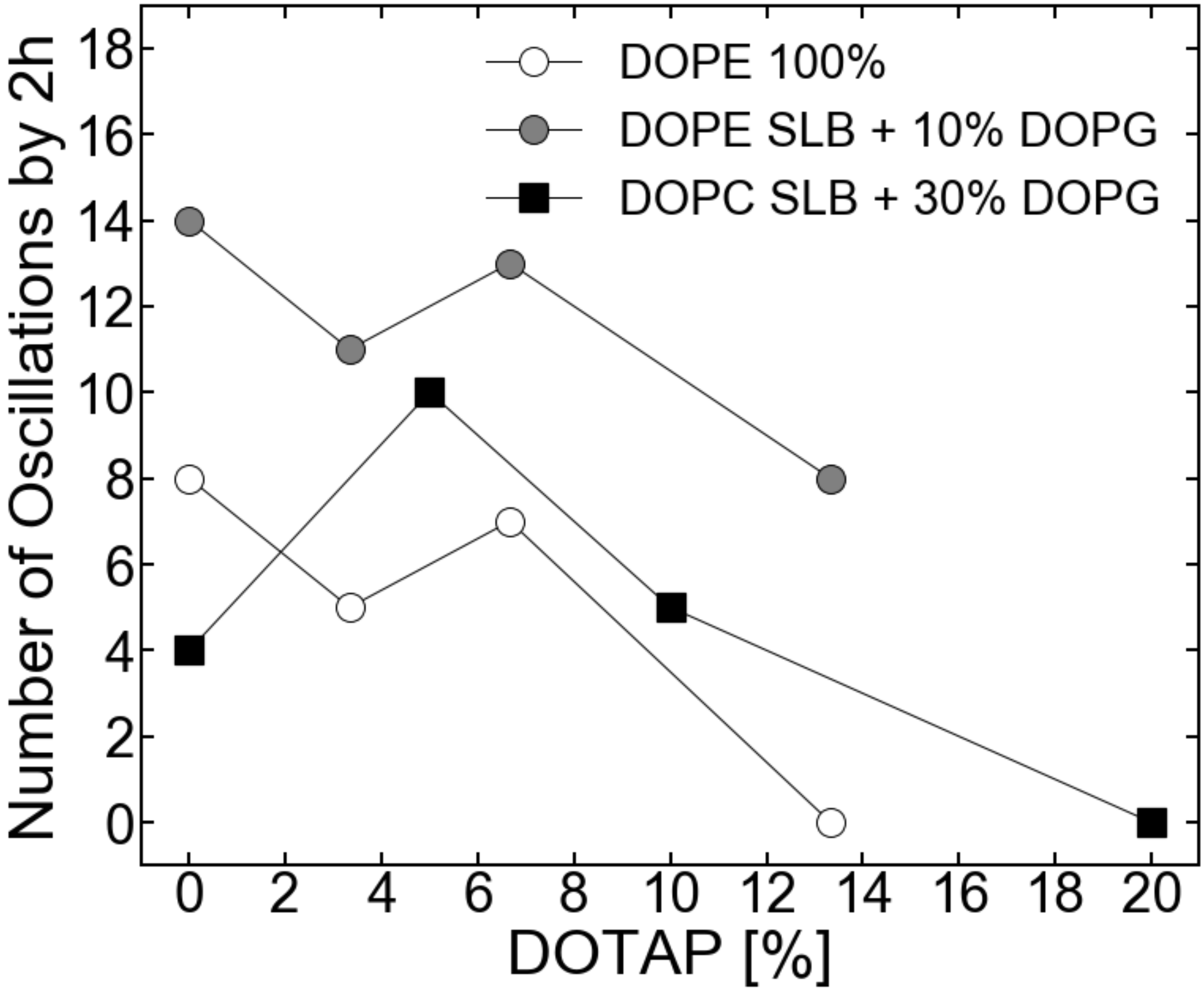


**Supplementary Figure 25.** The number of oscillations within the first 2 hours of MinD TXTL reactions (P70a-*minD*, 10 nM) as a function of the relative DOTAP concentration in a pure DOPE SLB, a DOPG/DOPE SLBs, and a DOPG/DOPC SLB.

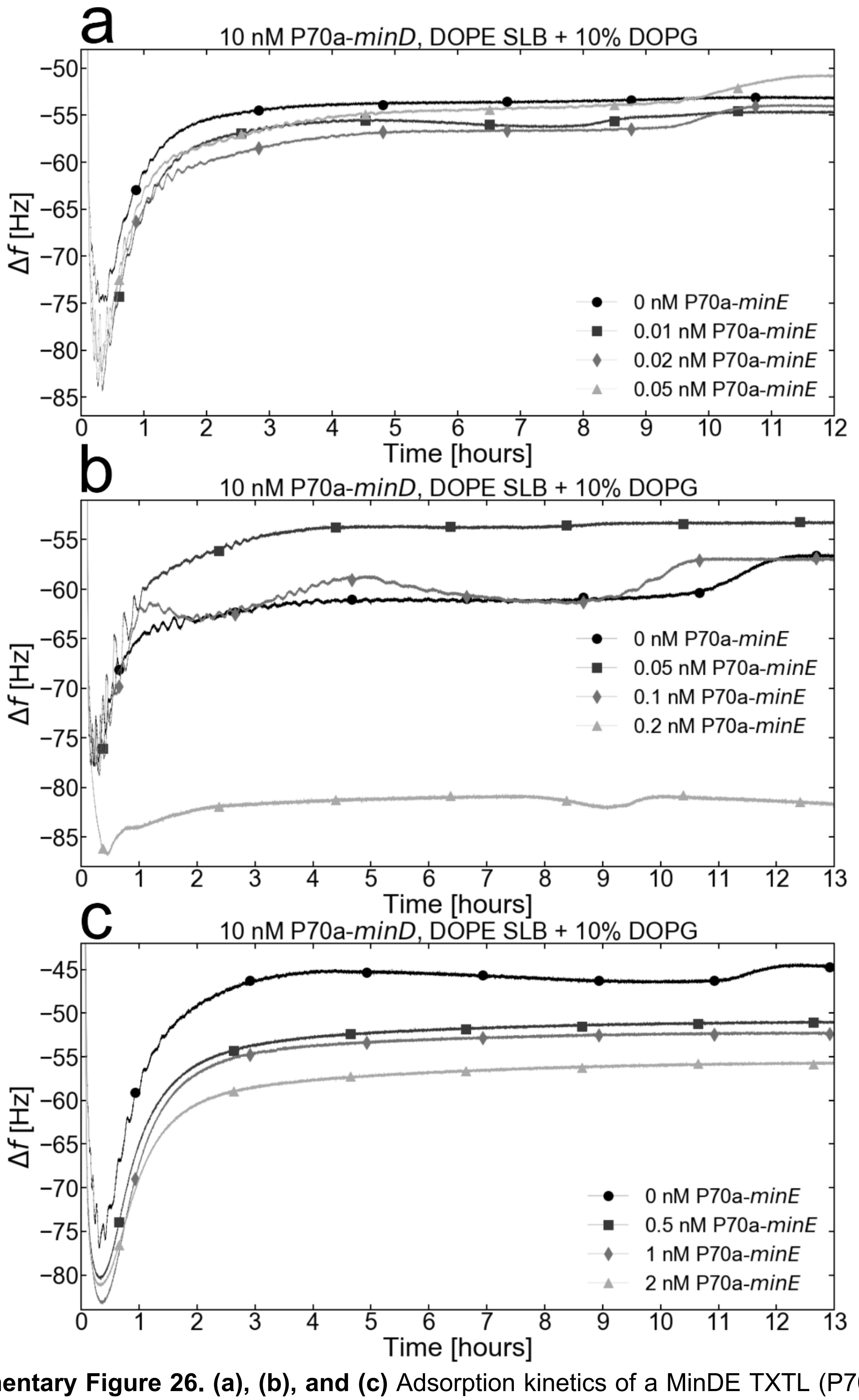


**Supplementary Figure 26. (a), (b), and (c)** Adsorption kinetics of a MinDE TXTL (P70a-*minD*, 10 nM, P70a-*minE*, varied) reaction for different P70a-*minE* concentrations into a DOPG/DOPE SLB.

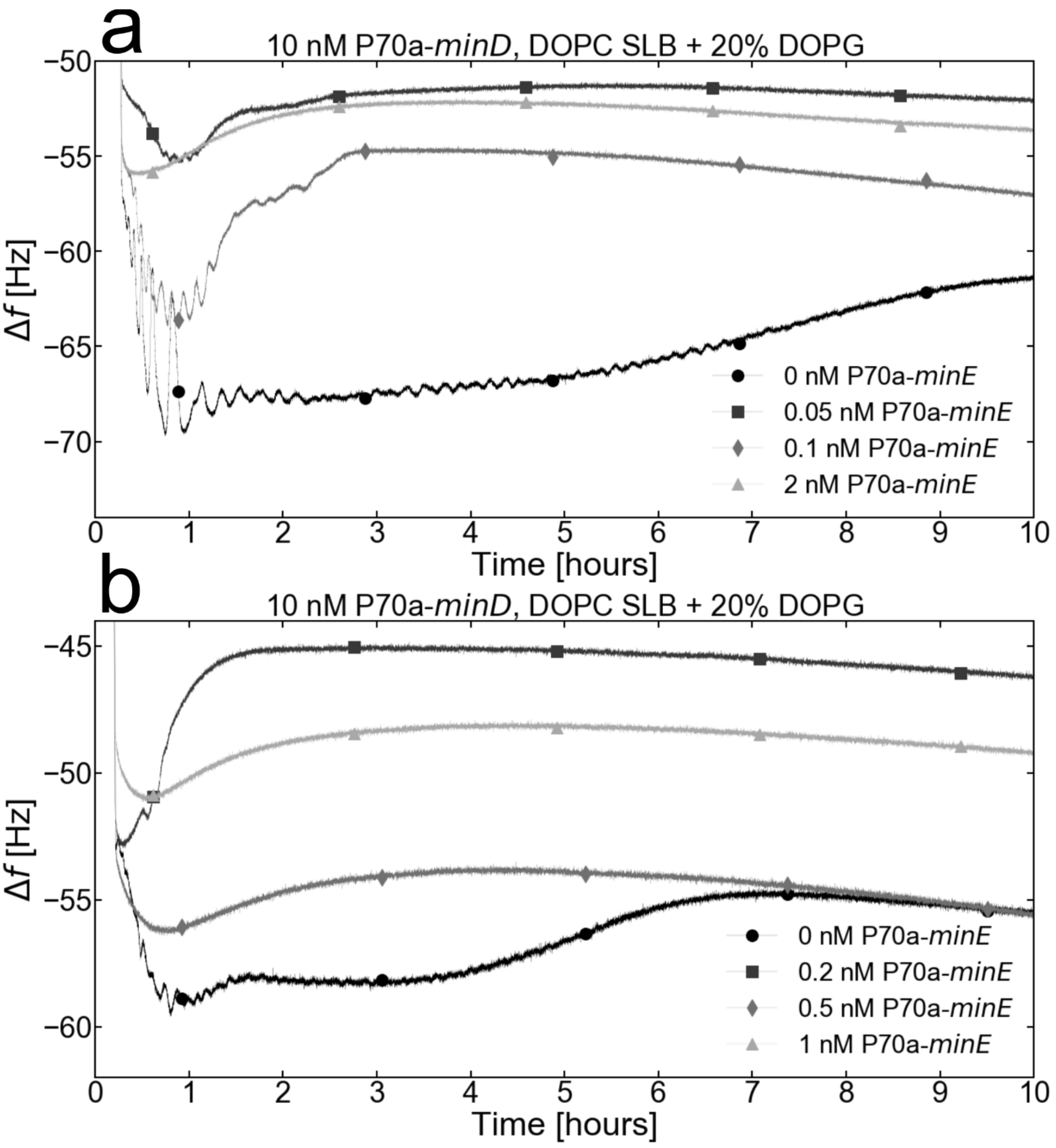


**Supplementary Figure 27. (a) and (b)** Adsorption kinetics of a MinDE TXTL (P70a-*minD*, 10 nM, P70a-*minE*, varied) reaction for different P70a-*minE* concentrations into a DOPG/DOPC SLB.

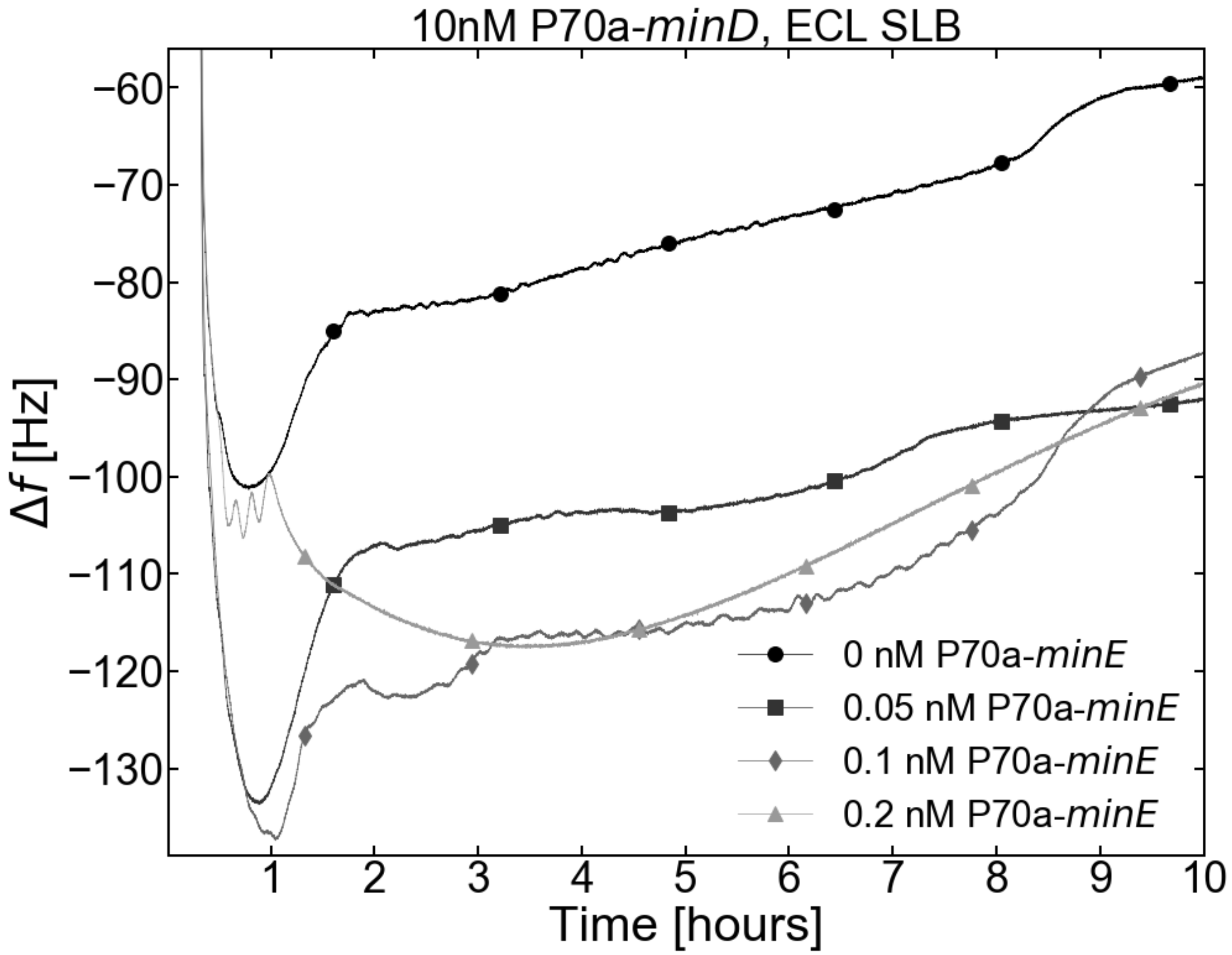


**Supplementary Figure 28.** Adsorption kinetics of a MinDE TXTL (P70a-*minD*, 10 nM, P70a-*minE*, varied) reaction for different P70a-*minE* concentrations into a ECL SLB.

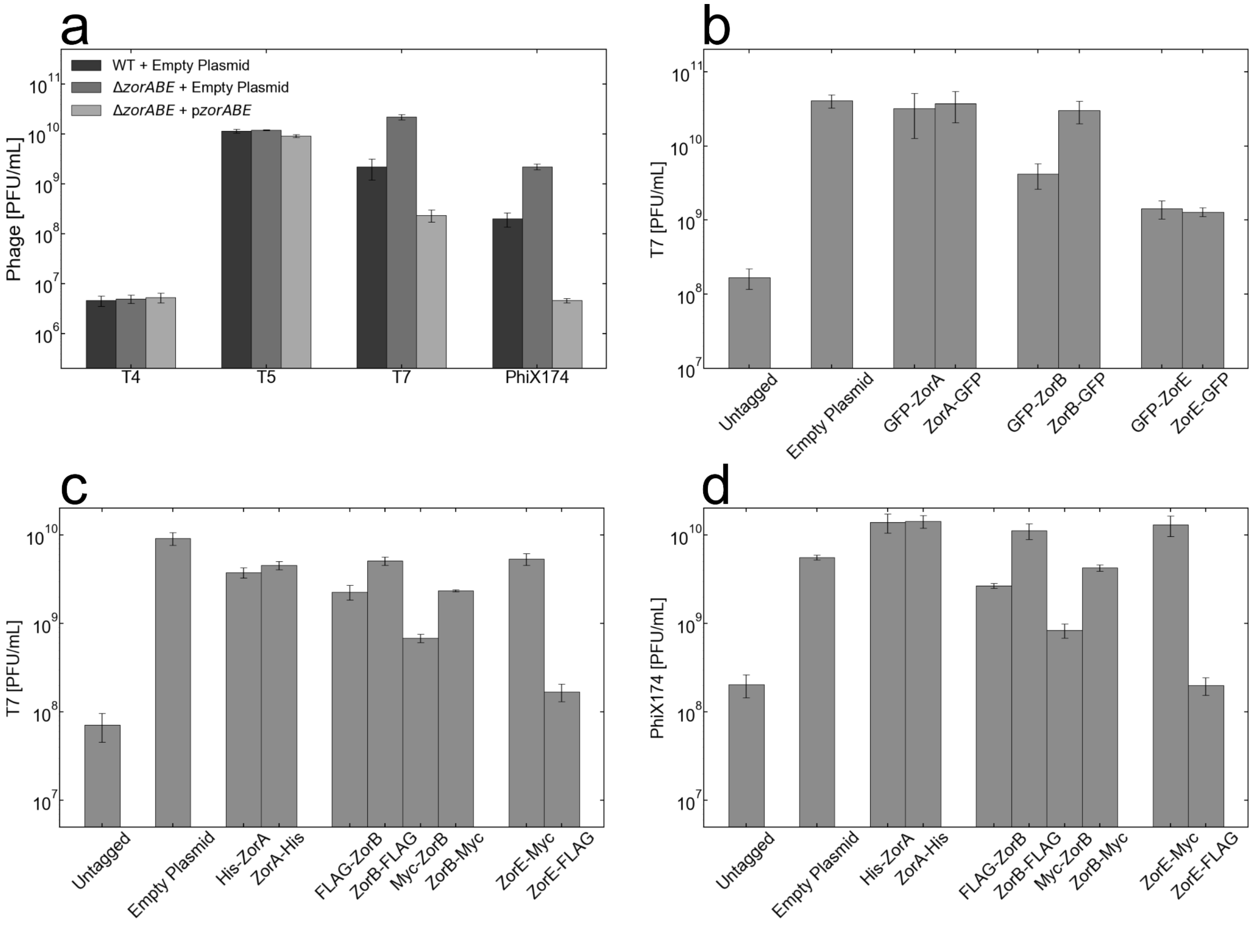


**Supplementary Figure 29. Tagging of Zorya proteins disrupts defense against phages. (a)** Reduction in plaque forming units in the presence of Zorya. **(b)** Reduction in T7 phage plaque forming units in the Δ*zorABE* knockout mutant. Zorya was supplemented on plasmids containing different combinations of the GFP tag at the termini of the respective proteins. **(c)** Reduction in the T7 plaque forming units in the Δ*zorABE* knockout mutant. Zorya was supplemented on plasmids containing different combinations of His, Myc, and FLAG tags at the termini of the respective proteins. **(d)** Reduction in the phiX174 plaque forming units in the Δ*zorABE* knockout mutant. Zorya was supplemented on plasmids containing different combinations of His, Myc, and FLAG tags at the termini of the respective proteins.

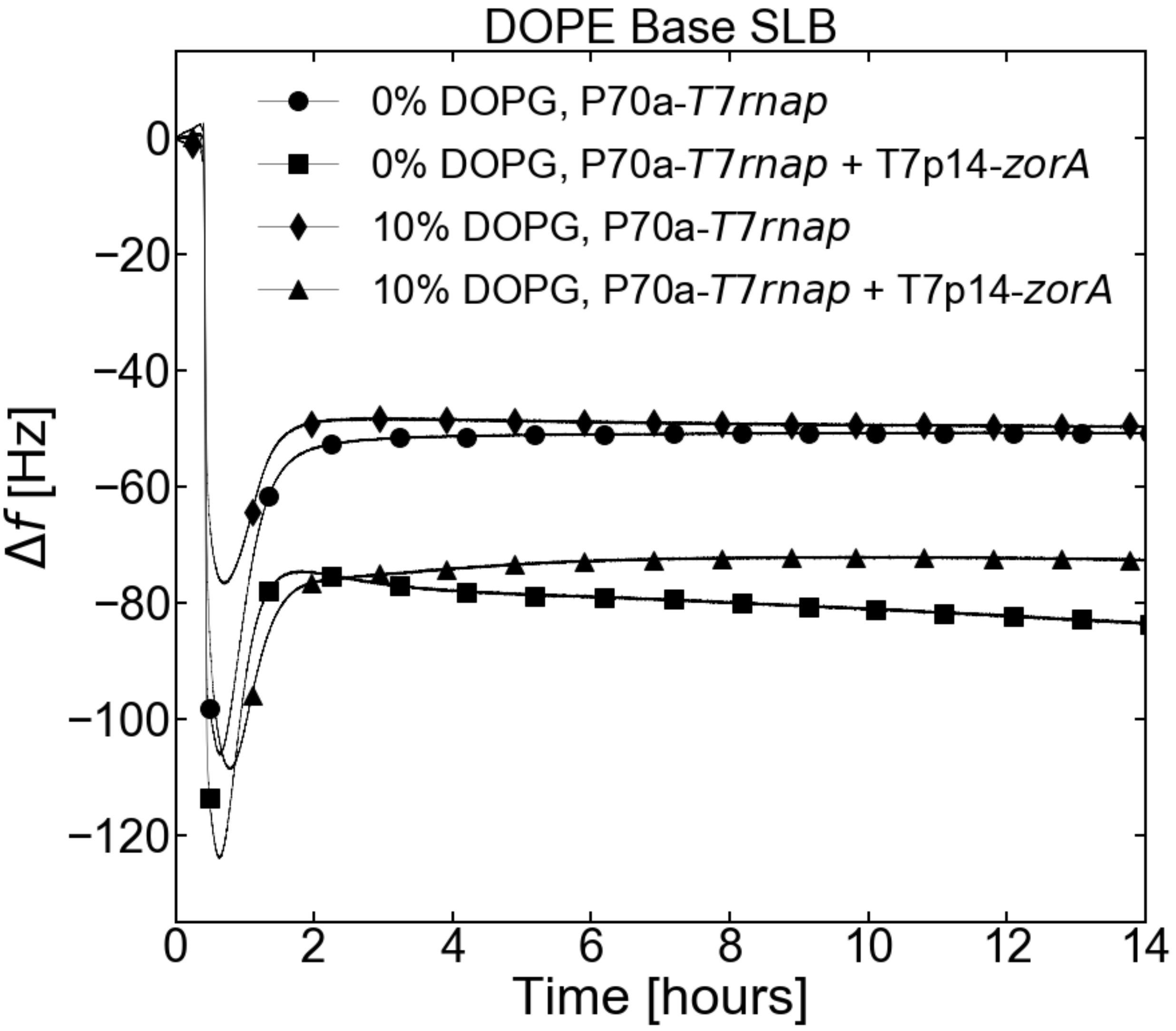


**Supplementary Figure 30.** Adsorption kinetics of a blank (P70a-*T7rnap*, 0.2 nM) and ZorA (P70a-*T7rnap*, 0.2 nM, T7p14-*zorA*, 10 nM) conditions with either a pure DOPE or DOPE + 10% DOPG SLBs.

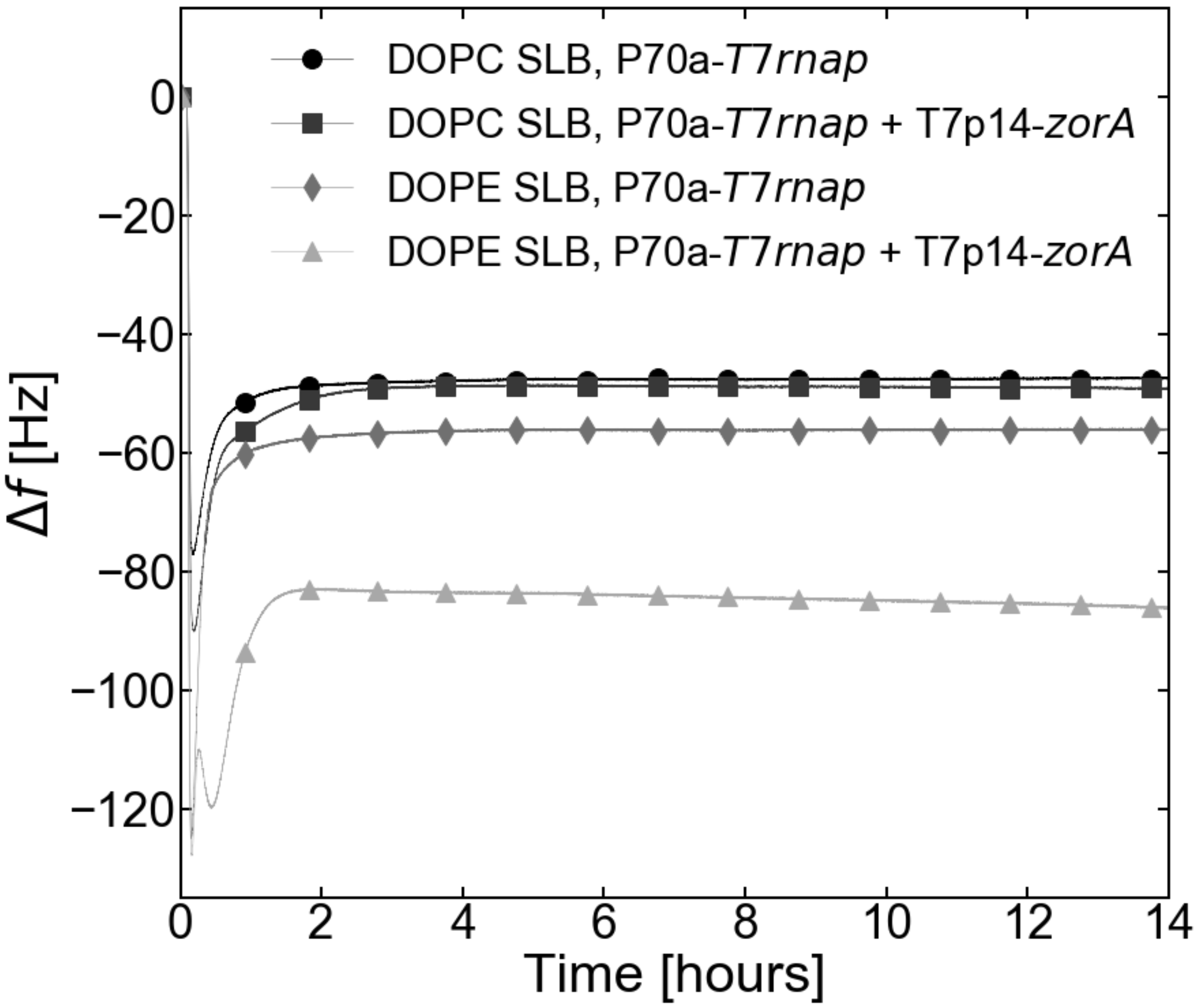


**Supplementary Figure 31.** The adsorption kinetics of a blank (P70a-*T7rnap*, 0.2 nM) and ZorA (P70a-*T7rnap*, 0.2 nM, T7p14-*zorA*, 10 nM) conditions with either a pure DOPC or pure DOPE SLBs.

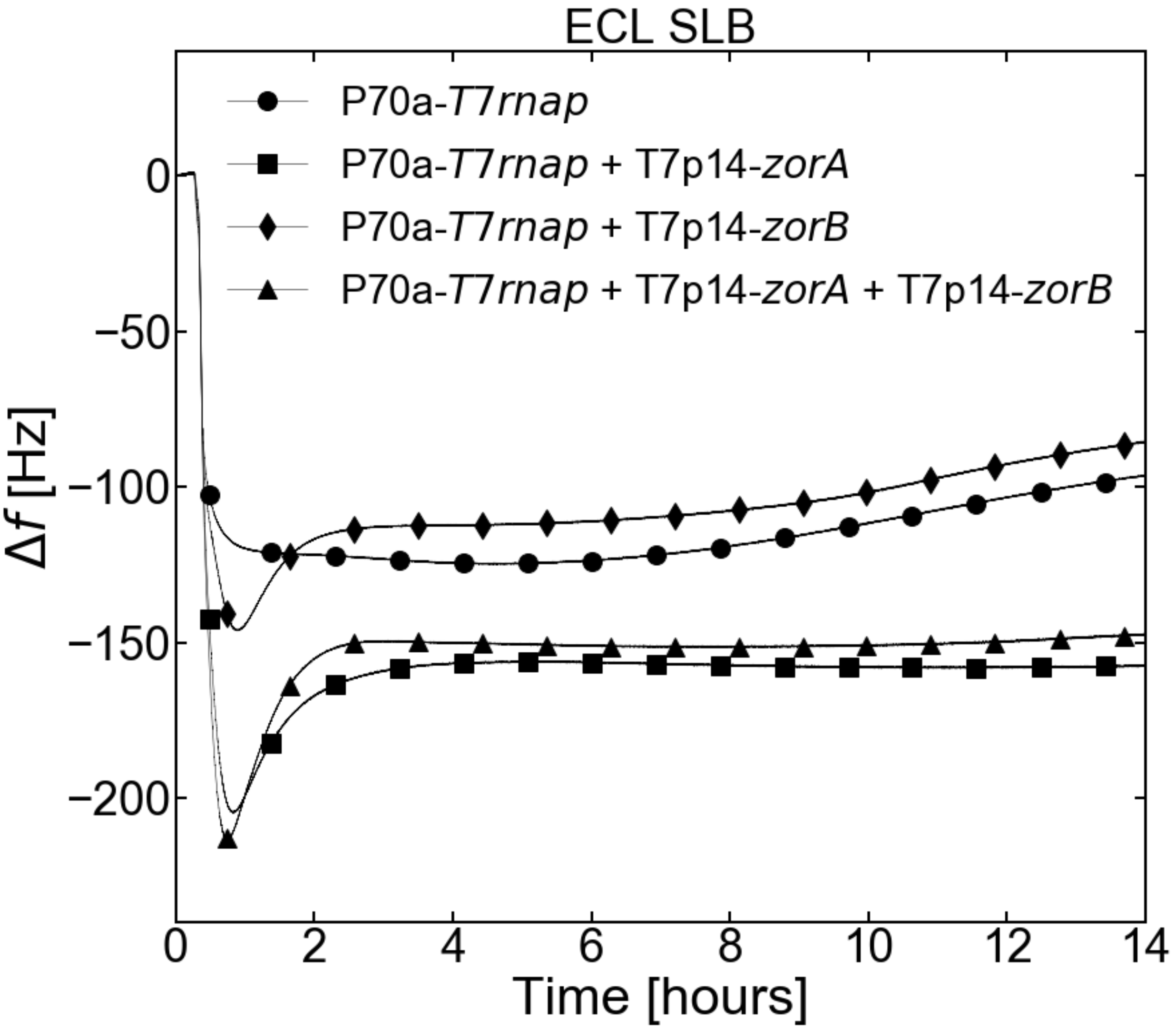


**Supplementary Figure 32.** The adsorption kinetics of a blank (P70a-*T7rnap*, 0.2 nM), ZorA alone (P70a-*T7rnap*, 0.2 nM, T7p14-*zorA*, 10 nM), ZorB alone (P70a-*T7rnap*, 0.2 nM, T7p14-*zorB*, 10 nM), and ZorA and ZorB together (P70a-*T7rnap*, 0.2 nM, T7p14-*zorA*, 10 nM, and T7p14-*zorB*, 10 nM) with an ECL SLB.

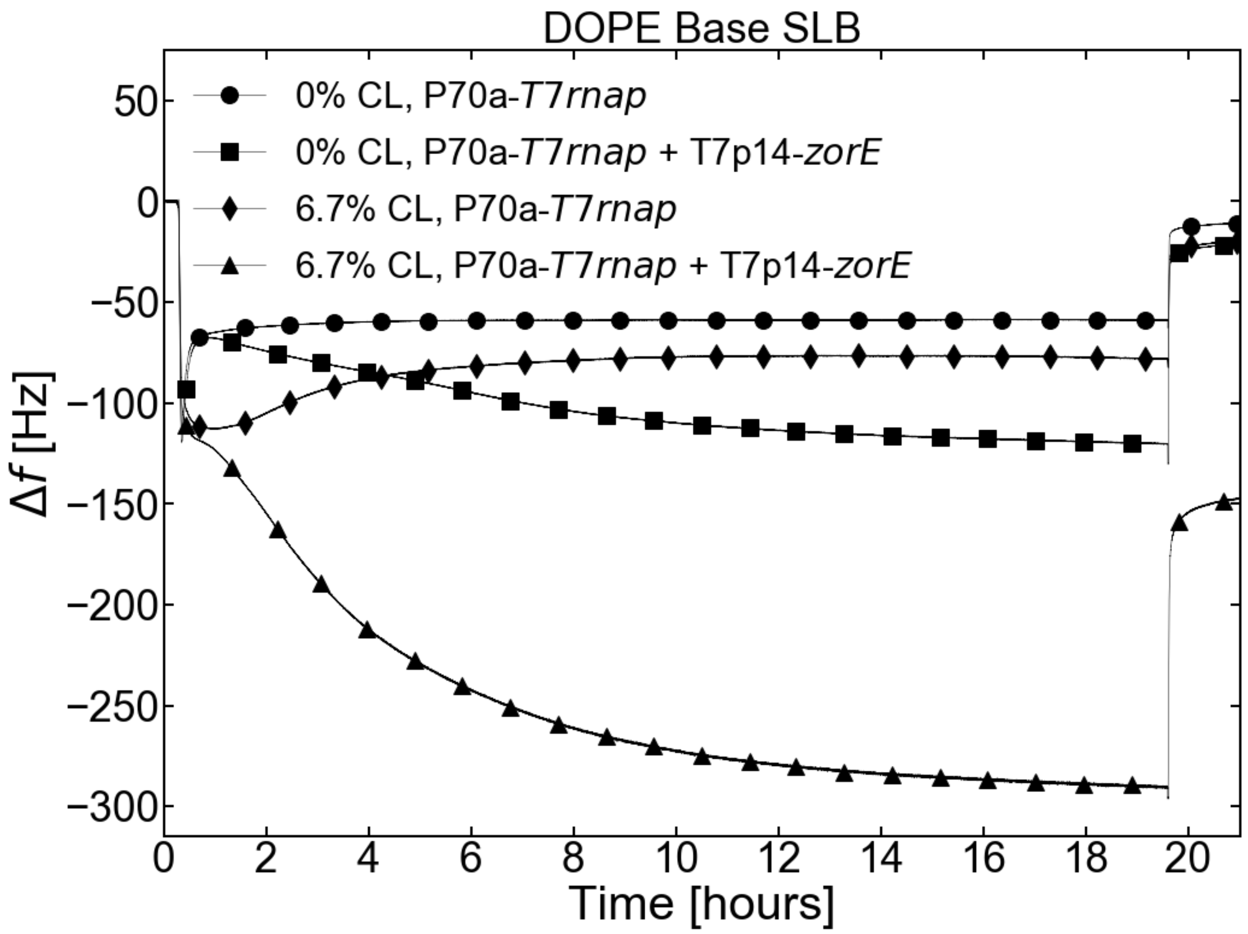


**Supplementary Figure 33.** Same as in **Fig. 6d** but with the measurement extended to t = 21 h after TXTL incubation start. The 1-h Tris NaCl flush starts at t = 19.6 h and the stabilized level is used for **Fig. 6e**.

| SLB composition | concentration (mM) | mass density (mg/mL) |
|---|---|---|
| DOPC | 1.3 mM | 1 mg/mL |
| EggPC | 1.3 mM | 1 mg/mL |
| DOPE | 1.95 mM | 1.5 mg/mL |
| *E. coli* lipids (ECL) | 3 mM (approximated) | 3 mg/mL |

**Supplementary Table S1**. Concentrations of phospholipids in IPA during SALB formation to obtain full coverage of the QCMD sensor and no nonspecific TXTL adsorption. Note that ECL are a mix of several phospholipids of different molecular weights, the average molecular weight of which is not communicated by the manufacturer but roughly estimated from its approximate composition.
